# Understanding and imaging PHGDH-driven intrinsic resistance to mutant IDH inhibition in gliomas

**DOI:** 10.1101/2025.08.30.673205

**Authors:** Céline Taglang, Georgios Batsios, Anne Marie Gillespie, Joanna J Phillips, Jennie W Taylor, Pavithra Viswanath

## Abstract

Mutations in isocitrate dehydrogenase (IDHm) define a distinct molecular class of gliomas. IDHm converts α-ketoglutarate (α-KG) to the oncometabolite D-2-hydroxyglutarate (D-2HG), which drives tumorigenesis. The IDHm inhibitor vorasidenib suppresses D-2HG production and extends progression-free survival in some, but not all, IDHm glioma patients. Here, using clinically relevant patient-derived IDHm models and patient tissue, we show that phosphoglycerate dehydrogenase (PHGDH) drives intrinsic resistance to vorasidenib by promiscuously converting α-KG to D-2HG and maintaining D-2HG concentration despite IDHm inhibition. Silencing PHGDH sensitizes resistant models to vorasidenib, while conversely, overexpressing PHGDH induces vorasidenib resistance in sensitive models. Importantly, deuterium metabolic imaging of D-2HG production from diethyl-[3,3’-^2^H]-α-ketoglutarate provides an early readout of response and resistance to vorasidenib that is not available by anatomical imaging *in vivo.* Collectively, we have identified PHGDH-driven D-2HG production as an intrinsic mechanism of resistance to vorasidenib and diethyl-[3,3’-^2^H]α-ketoglutarate as a non-invasive tracer for interrogating intrinsic resistance in IDHm gliomas.

**STATEMENT OF SIGNIFICANCE:** Vorasidenib, which suppresses D-2HG production, is the first precision therapy to be approved for IDHm glioma patients. We show that PHGDH-driven restoration of D-2HG production mediates intrinsic resistance to vorasidenib in IDHm gliomas. Importantly, deuterium metabolic imaging of D-2HG production from diethyl-[3,3’-^2^H]-α-ketoglutarate enables non-invasive assessment of resistance in IDHm gliomas.

## INTRODUCTION

Gliomas are incurable, malignant primary brain tumors in adults (1). The discovery of driver mutations in isocitrate dehydrogenase 1 or, less commonly, 2 (IDHm) has revolutionized glioma classification by introducing molecular classes with prognostic and therapeutic implications (2,3). Tumors with wild-type isocitrate dehydrogenase (IDHwt) are classified as glioblastoma (GBM, grade 4), while IDHm tumors are further classified as oligodendrogliomas (IDHm in combination with a 1p19q codeletion, grades 2-3) or astrocytomas (IDH, 1p/19q intact, grades 2-4) (2,3). Although IDHm gliomas initially grow slowly compared to IDHwt tumors and have a better prognosis, they tend to afflict younger patients (median age of ∼40 years for IDHm vs. ∼70 years for IDHwt), who are likely to be socially and economically active (4,5). Standard of care for IDHm glioma patients includes maximal safe surgical resection, radiation, and/or chemotherapy (3). However, tumor recurrence is inevitable, and both radiation and chemotherapy are associated with adverse effects on physical and cognitive abilities, which is particularly devastating to this relatively young patient population (3,6,7).

The IDHwt enzyme converts isocitrate to α-ketoglutarate (α-KG), which serves as a cofactor for >60 dioxygenases, including the TET family of DNA 5-methylcytosine (5mC) hydroxylases, the KDM family of histone demethylases, and prolyl hydroxylases (8). In contrast, point mutations in the active site of the IDHm enzyme, most commonly IDH1 R132H, lead to the neomorphic ability to convert α-KG to the oncometabolite D-2-hydroxyglutarate (D-2HG) (8,9). D-2HG competitively inhibits the activity of α-KG-dependent enzymes, leading to hypermethylation of histones and DNA and a block in differentiation that drives tumorigenesis (8,9).

IDHm is an attractive therapeutic target because it is the earliest known genetic alteration in these tumors and is largely retained at recurrence (3,8,9). Studies with doxycycline-inducible IDHm expression suggest that the effects of D-2HG on histone and DNA hypermethylation and differentiation are, at least, partially reversible (10). Several IDHm inhibitors, such as vorasidenib (VOR; brain-penetrant dual inhibitor of mutant IDH1 and IDH2), ivosidenib (IVO; inhibitor of mutant IDH1), safusidenib, and olutasidenib (brain-penetrant inhibitors of mutant IDH1), have been developed and shown to suppress D-2HG production and delay tumor growth in mice bearing flank or intracranial IDHm xenografts (11–15). Early phase clinical trials with these drugs indicated that patients with contrast-enhancing tumors responded poorly to IDHm inhibition, with the majority of patients experiencing disease progression (16–18). In contrast, patients with non-contrast-enhancing tumors fared better, with the majority of patients experiencing stable disease (16–18). Importantly, in the phase 3 INDIGO trial, VOR significantly extended progression-free survival (27.7 months) relative to a placebo (11.1 months) in patients with grade 2 IDHm tumors who received no other prior treatment (besides surgery) (19). Based on these results, VOR has been approved for the treatment of IDHm glioma patients. While undoubtedly significant, the 28% progression rate reported in VOR-treated patients in the INDIGO trial and the observation that disease progression occurred within ∼18 months point to the existence of intrinsic or rapidly acquired secondary resistance to VOR in IDHm glioma patients.

Magnetic resonance imaging (MRI) is the mainstay for glioma patient management (20,21). However, IDHm gliomas are difficult to visualize because they typically do not enhance on post-gadolinium MRI (20). Importantly, assessing response to therapy in IDHm glioma patients using MRI is challenging (22,23). Of note, IDHm inhibitors such as VOR often have a cytostatic effect of delaying tumor growth rather than inducing radiographically observable tumor regression via cytotoxic cell death, which emphasizes the importance of measuring response using a parameter other than MRI-detectable tumor size (23–25). Since D-2HG accumulates to millimolar concentrations (5-35 mM) in IDHm gliomas and is virtually undetectable in the normal brain and IDHwt GBM (8), imaging D-2HG production has the potential to enable non-invasive assessment of metabolically active tumor tissue. Magnetic resonance spectroscopy (MRS) is an MR-compatible method of imaging tissue metabolites with concentrations >1 mM (26). ^1^H-MRS interrogates the magnetic resonance of protons in tissue metabolites and provides a readout of steady-state metabolite abundances (26). ^13^C-MRS, following the administration of a ^13^C-labeled substrate, is the gold standard for measuring the activity of a metabolic pathway (26). However, the sensitivity of ^13^C-MRS is not sufficient for non-invasive imaging *in vivo.* Deuterium metabolic imaging (DMI) interrogates the magnetic resonance of deuterons (^2^H) and can be used to trace the metabolic fate of ^2^H-labeled substrates in live cells, animals, and patients (27). DMI, following the administration of [6,6-^2^H]-glucose, has been used to monitor the Warburg effect in preclinical glioma models and adult GBM patients (27,28). However, whether DMI can be used to monitor α-KG metabolism in IDHm gliomas is unknown.

The goal of this study was to interrogate the molecular mechanisms driving resistance to VOR and to develop a DMI-based tracer for imaging response and resistance to VOR in preclinical IDHm glioma models. Using a panel of 12 clinically relevant patient-derived IDHm glioma models and glioma patient tissue, we show that intrinsic resistance is driven by restoration of D-2HG synthesis despite treatment with VOR. Mechanistically, phosphoglycerate dehydrogenase (PHGDH), which promiscuously converts α-KG to D-2HG, drives VOR resistance and represents a collateral vulnerability in the context of IDHm inhibition. Importantly, we demonstrate that DMI following the administration of diethyl-[3,3’-^2^H]-α-KG informs on early response to therapy and intrinsic resistance to VOR in IDHm gliomas.

## RESULTS

### Intrinsic resistance to VOR is observed in patient-derived IDHm glioma models

First, we examined the response of a panel of 12 patient-derived IDHm glioma models to VOR. These models were isolated from patients with grade 2-4 astrocytomas or oligodendrogliomas harboring endogenous IDHm and various secondary and tertiary genetic alterations (Supplementary Fig. S1A). All of these models have previously been described (29–36) and were grown as neurospheres in culture. Using an enzymatic assay that specifically quantifies D-2HG (37), we confirmed that IDHm neurospheres produced high concentrations of D-2HG (0.5-15 mM), in contrast to primary human astrocytes and a patient-derived GBM model (GBM6), which produced <0.1 mM D-2HG (Supplementary Fig. S1B). We then quantified the IC50 for VOR by measuring the effect of various doses of VOR on the clonogenicity of patient-derived IDHm glioma neurospheres. As shown in Fig. 1A, 6 models were sensitive to VOR with IC50 values <0.3 μM (astrocytoma: BT257, BT142, SF10602; oligodendroglioma: SF10417, BT54, TS603) while the other 6 models were resistant with IC50s >5 μM (astrocytoma: MGG119, MGG152, NCH1681, TB096, GSC580; oligodendroglioma: NCH612). We orthogonally confirmed these results by quantifying the effect of treatment with VOR at IC50 on the expression of Ki67, which is a marker of proliferation (Fig. 1B). Although VOR did not induce cell death in sensitive or resistant models, it induced cellular differentiation as measured by the expression of the normal astrocytic marker GFAP in sensitive but not resistant models (Supplementary Fig. S1C-S1D). Furthermore, we observed a similar pattern of intrinsic resistance to IVO in our IDHm glioma models (Supplementary Fig. S1E). Importantly, given that all of the resistant IDHm glioma models were established from patients who had never been treated with an IDHm inhibitor (29–36), these results point to the existence of intrinsic resistance to IDHm inhibition in gliomas.

**Figure 1.**
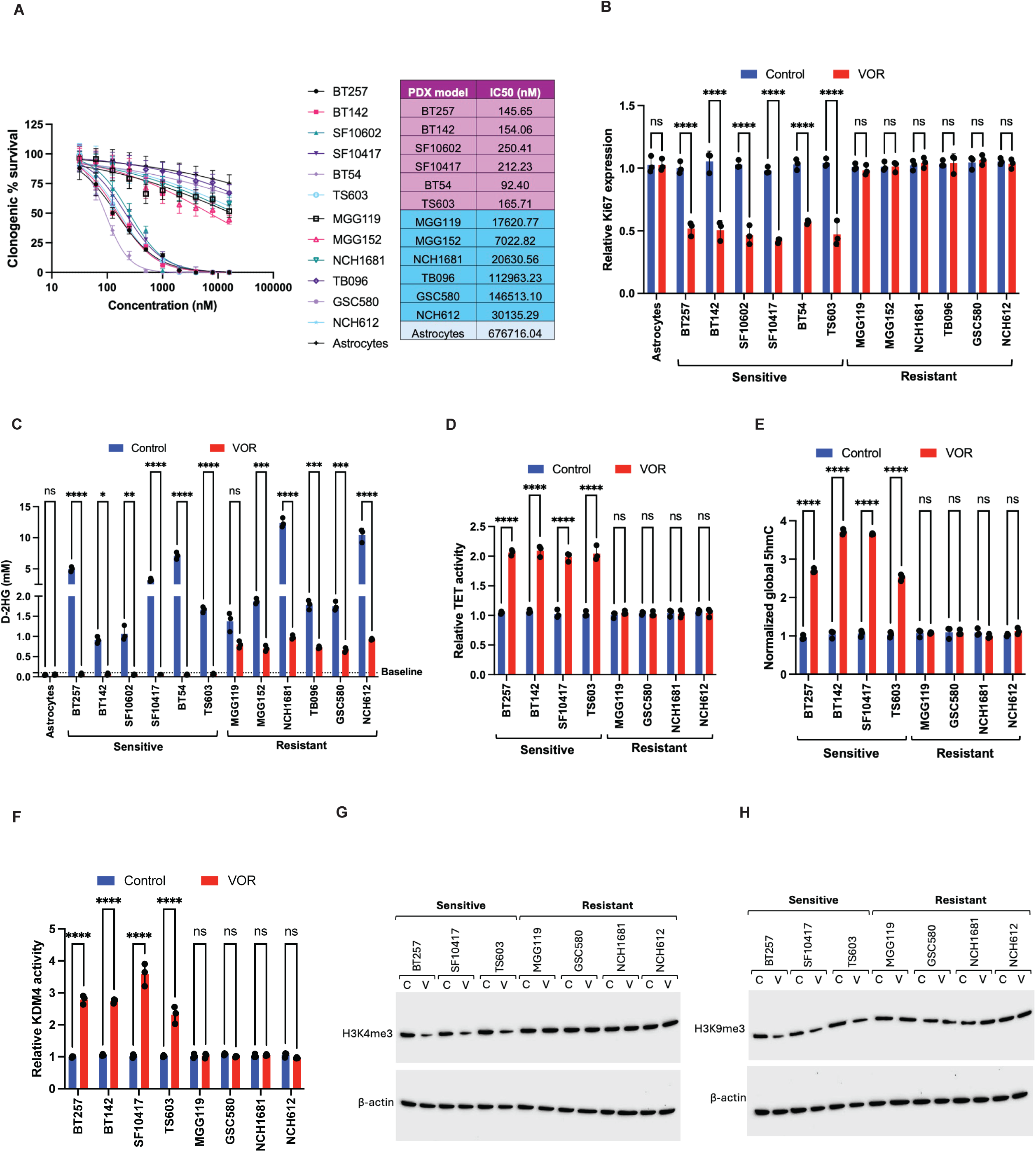
Intrinsic resistance to VOR is observed in patient-derived IDHm glioma models. Effect of VOR on clonogenicity **(A)**, Ki67 expression **(B)**, D-2HG concentration **(C)**, TET DNA 5mC hydroxylase activity **(D)**, global 5hMC abundance **(E)**, and KDM4 demethylase activity **(F)** of patient-derived IDHm glioma models. Western blots showing the effect of VOR on global H3K4me3 **(G)** and H3K9me3 **(H)** in patient-derived IDHm glioma models. All studies were n≥3. Data is presented as mean ± standard deviation. * indicates statistical significance with p<0.05, ** indicates p<0.01, *** indicates p<0.001, **** indicates p<0.0001, ns is not significant.

### Restoration of D-2HG production facilitates intrinsic resistance to VOR in patient-derived IDHm glioma models

Next, we quantified the effect of VOR on D-2HG production in our models. As shown in Fig. 1C, VOR reduced D-2HG abundance in sensitive models to levels observed in vehicle-treated controls and astrocytes (<0.1 mM). In contrast, resistant models retained significant abundance of D-2HG (>0.6 mM) despite treatment with VOR (Fig. 1C). Since D-2HG competitively inhibits α-KG-dependent enzymes such as the KDM histone demethylases and TET DNA 5-mC hydroxylases with IC50 values of ∼0.2 mM (38,39), D-2HG concentrations of >0.6 mM are expected to be sufficient to block histone and DNA demethylation in resistant models. Indeed, as shown in Fig. 1D-1E, VOR significantly increased TET enzyme activity and DNA hydroxylation (the first step in demethylation) in sensitive but not resistant models. Similarly, VOR significantly increased KDM4A demethylase activity and reduced global trimethylation of histone H3K4 (H3K4me3) and H3K9 (H3K9me3) in sensitive but not resistant IDHm glioma models (Fig. 1F-1H).

To confirm that intrinsic resistance to VOR is associated with restoration of D-2HG synthesis, we used ^13^C-MRS to trace D-2HG production from [1-^13^C]-α-KG in our IDHm glioma models. We examined primary human astrocytes as baseline controls. Metabolism of [1-^13^C]-α-KG (170.6 ppm) via IDHm will yield [1-^13^C]-D-2HG (183.2 ppm) while transamination mediated by alanine aminotransferases (GPT1, GPT2), aspartate aminotransferases (GOT1, GOT2), branched chain aminotransferases (BCAT1, BCAT2), or glutamate dehydrogenases (GLUD1, GLUD2) will produce [1-^13^C]-glutamate (175.1 ppm). As shown in Fig. 2A-2C and Supplementary Fig. S1F, astrocytes metabolized [1-^13^C]-α-KG to [1-^13^C]-glutamate, and vehicle-treated IDHm glioma cells shunted [1-^13^C]-α-KG to [1-^13^C]-D-2HG and reduced metabolism to [1-^13^C]-glutamate. Importantly, treatment with VOR normalized [1-^13^C]-D-2HG and [1-^13^C]-glutamate to baseline astrocyte levels in VOR-sensitive IDHm models (Fig. 2A-2C and Supplementary Fig. S1F). In contrast, VOR-resistant models retained significant synthesis of [1-^13^C]-D-2HG despite treatment with VOR (Fig. 2A-2C and Supplementary Fig. S1F). VOR did not alter [1-^13^C]-glutamate concentration in VOR-resistant models (Supplementary Fig. S1F). Collectively, these results indicate that restoration of D-2HG synthesis from α-KG is a potential mechanism of intrinsic resistance to VOR in IDHm glioma cells.

**Figure 2.**
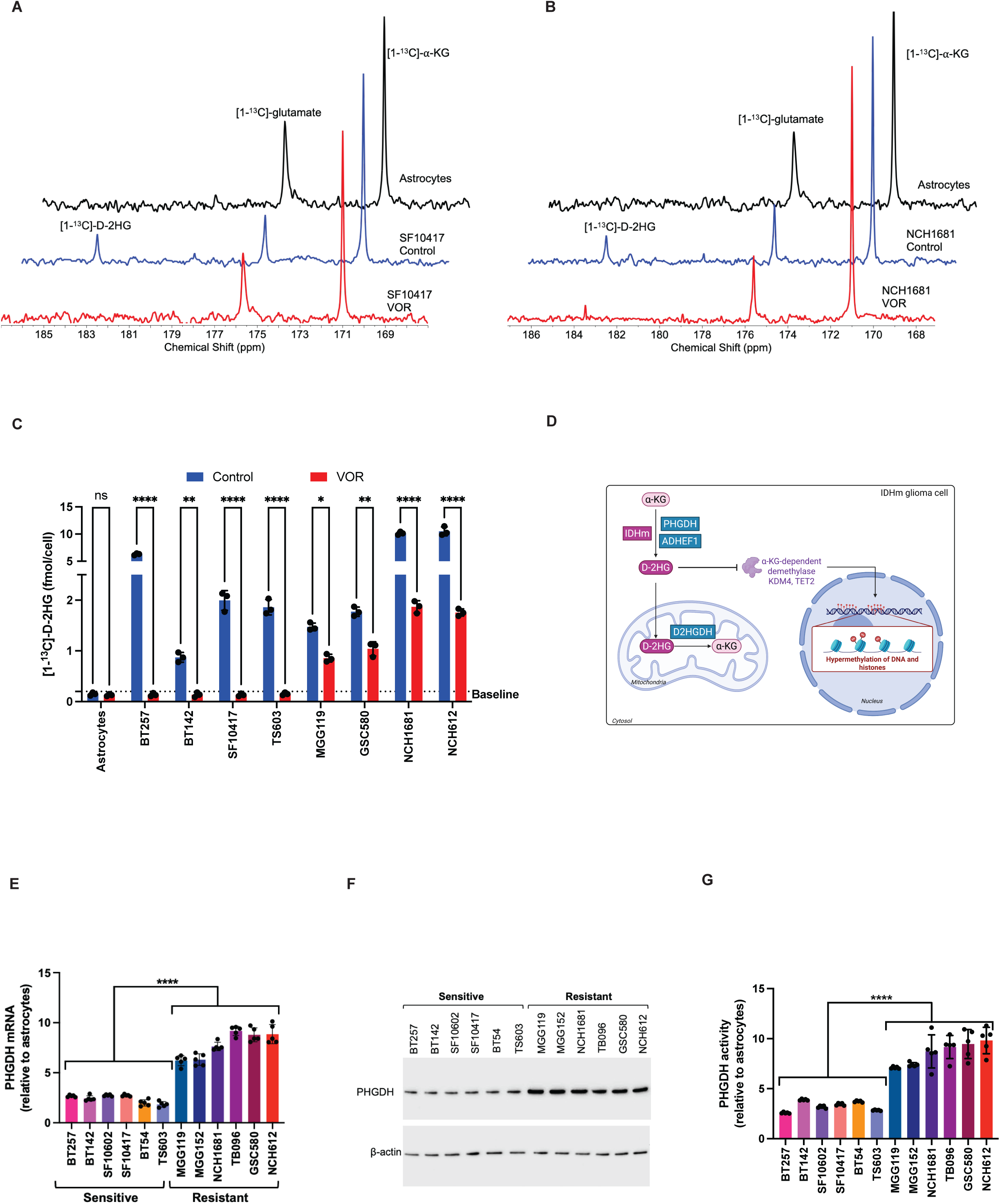
Restoration of D-2HG synthesis from α-KG mediates intrinsic resistance to VOR in IDHm gliomas. **(A)** Representative ^13^C-MR spectra of [1-^13^C]-α-KG metabolism in VOR-sensitive SF10417 cells treated with vehicle or VOR for 72 h. Primary human astrocytes were used as baseline controls. **(B)** Representative ^13^C-MR spectra of [1-^13^C]-α-KG metabolism in VOR-resistant NCH1681 cells treated with vehicle or VOR for 72 h. Primary human astrocytes were used as baseline controls. **(C)** Quantification of D-2HG production from [1-^13^C]-α-KG in VOR-sensitive (BT257, BT142, SF10417, TS603) and VOR-resistant (MGG119, GSC580, NCH1681, and NCH612) IDHm glioma cells. Primary human astrocytes were used as baseline controls for D-2HG concentration, as indicated by the dotted line. **(D)** Schematic illustration of the metabolic determinants of D-2HG abundance in mammalian cells. PHGDH mRNA **(E)**, PHGDH protein **(F)**, and PHGDH activity **(G)** in VOR-sensitive and VOR-resistant IDH glioma neurospheres. All studies were n≥3. Data is presented as mean ± standard deviation. * indicates statistical significance with p<0.05, ** indicates p<0.01, *** indicates p<0.001, **** indicates p<0.0001, ns is not significant.

### Phosphoglycerate dehydrogenase (PHGDH) is upregulated in patient-derived IDHm glioma cells that are intrinsically resistant to VOR

Next, we examined the mechanism by which D-2HG synthesis is restored in IDHm gliomas with intrinsic resistance to VOR. α-KG can be converted to D-2HG in cells lacking IDHm via the promiscuous activity of cellular dehydrogenases such as PHGDH or hydroxyl acid-oxoacid-transhydrogenase (ADHFE1) (40,41). Conversely, loss of D-2HG dehydrogenase (D2HGDH), the enzyme that normally degrades D-2HG, can also lead to the accumulation of D-2HG (40) (see Fig. 2D). As shown in Fig. 2E-2G, PHGDH mRNA, protein, and activity were significantly higher in VOR-resistant models relative to sensitive models at baseline (in the absence of VOR treatment). In contrast, we did not observe consistent changes in the expression of ADHFE1 or D2HGDH between VOR-sensitive and resistant neurospheres (Supplementary Fig. S2A-S2B).

### PHGDH drives D-2HG production from α-KG and induces resistance to VOR in patient-derived IDHm glioma cells

To assess whether PHGDH drives D-2HG synthesis, we examined the effect of silencing PHGDH in VOR-resistant IDHm neurospheres (MGG119, GSC580, NCH1681, NCH612; Supplementary Fig. S2C). As shown in Fig. 3A-3B and Supplementary Fig. S2D, silencing PHGDH in combination with VOR in VOR-resistant neurospheres reduced [1-^13^C]-α-KG metabolism to [1-^13^C]-D-2HG and increased conversion to [1-^13^C]-glutamate to baseline levels similar to those observed in primary human astrocytes. We also confirmed that PHGDH silencing in combination with VOR reduced the steady-state pool of D-2HG to baseline in VOR-resistant neurospheres (Supplementary Fig. S2E). Importantly, PHGDH silencing significantly increased sensitivity to VOR (Fig. 3C). Furthermore, combined VOR and PHGDH silencing significantly increased KDM4 and TET activity and induced astrocytic differentiation in VOR-resistant IDHm glioma cells (Fig. 3D-3F).

**Figure 3.**
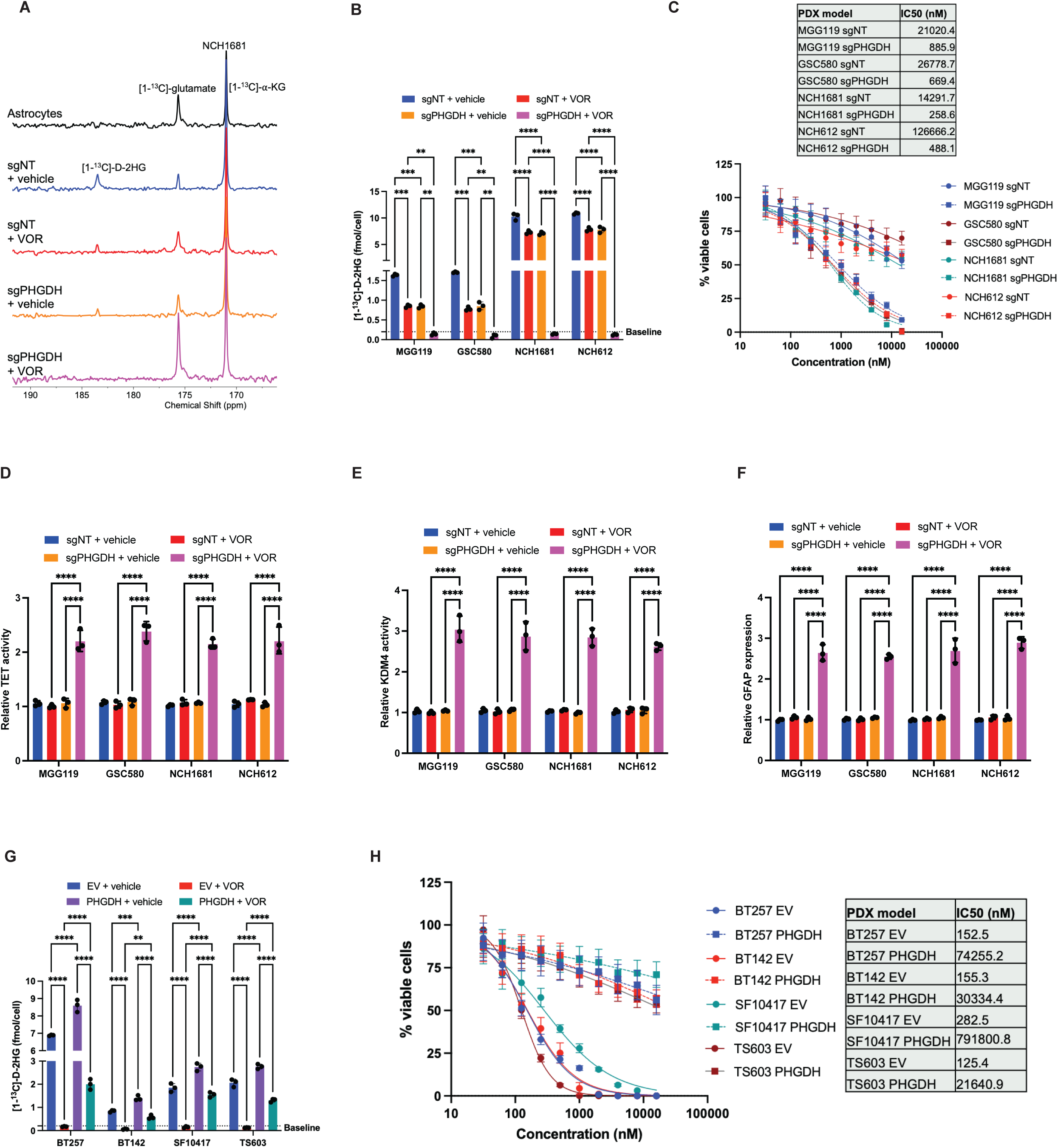
PHGDH drives D-2HG synthesis from α-KG in patient-derived IDHm glioma cells. Representative ^13^C-MR spectra of [1-^13^C]-α-KG metabolism **(A)** and quantification of ^13^C-labeled D-2HG synthesis **(B)** in NCH1681 cells expressing non-targeted sgRNA (sgNT) or PHGDH-targeted sgRNA (sgPHGDH). Cells were treated with vehicle or VOR for 72 h. **(C)** Effect of silencing PHGDH on the IC50 for VOR in VOR-resistant IDHm glioma cells (MGG119, GSC580, NCH1681, and NCH612). IC50 was measured using the clonogenicity assay. Effect of silencing PHGDH on TET DNA 5mC hydroxylase activity **(D)**, KDM4 activity **(E)**, and GFAP expression **(F)** in VOR-resistant IDHm glioma cells (MGG119, GSC580, NCH1681, and NCH612) treated with vehicle or VOR. **(G)** Quantification of ^13^C-labeled D-2HG synthesis from [1-^13^C]-α-KG in VOR-sensitive IDHm glioma models (BT257, BT142, SF10417, TS603) expressing an empty vector (EV) or PHGDH. Baseline D-2HG abundance is defined as the level in primary human astrocytes and is indicated by a dotted line. **(H)** Effect of overexpressing PHGDH on the sensitivity of IDHm glioma cells to VOR. All studies were n≥3. Data is presented as mean ± standard deviation. * indicates statistical significance with p<0.05, ** indicates p<0.01, *** indicates p<0.001, **** indicates p<0.0001, ns is not significant.

We also examined the effect of overexpressing PHGDH on IDHm gliomas that are intrinsically sensitive to VOR (astrocytoma: BT257, BT142; oligodendroglioma: SF10417, TS603; Supplementary Fig. S2F). As shown in Fig. 3G and Supplementary Fig. S2G, VOR reduced the synthesis of [1-^13^C]-D-2HG and increased the synthesis of [1-^13^C]-glutamate from [1-^13^C]-α-KG in cells expressing an empty vector, an effect that was abrogated by overexpression of PHGDH. Importantly, overexpressing PHGDH induced VOR resistance in patient-derived IDHm glioma cells that were intrinsically sensitive to VOR (Fig. 3H). Taken together, these results identify a role for PHGDH in driving D-2HG synthesis and, thereby, facilitating intrinsic resistance to IDHm inhibition in patient-derived IDHm glioma cells.

### PHGDH drives D-2HG production from α-KG and induces resistance to VOR in genetically engineered murine glioma cells and tumor xenografts

Patient-derived IDHm glioma models differ in secondary and tertiary genetic alterations besides IDHm that are critical for tumor growth (33). To validate the role of PHGDH in driving resistance to IDHm inhibition in an isogenic setting, we examined the effect of overexpressing PHGDH in a genetically engineered murine IDHm glioma model (NPAmut) (42). These cells were generated by Sleeping Beauty transposon-based activation of NRAS G12V and loss of TP53 and ATRX in combination with IDHm and have been extensively used to study the role of IDHm in glioma biology (42–45). Treatment with VOR suppressed [1-^13^C]-D-2HG synthesis in NPAmut cells expressing an empty vector (NPAmut_EV_; Fig. 4A-4B). In contrast, NPAmut cells overexpressing PHGDH (NPAmut_PHGDH_) retained significant labeling of [1-^13^C]-D-2HG from [1-^13^C]-α-KG despite treatment with VOR (Fig. 4A-4B). Importantly, overexpressing PHGDH increased the IC50 for VOR by > 100-fold in NPAmut cells (Fig. 4C).

**Figure 4.**
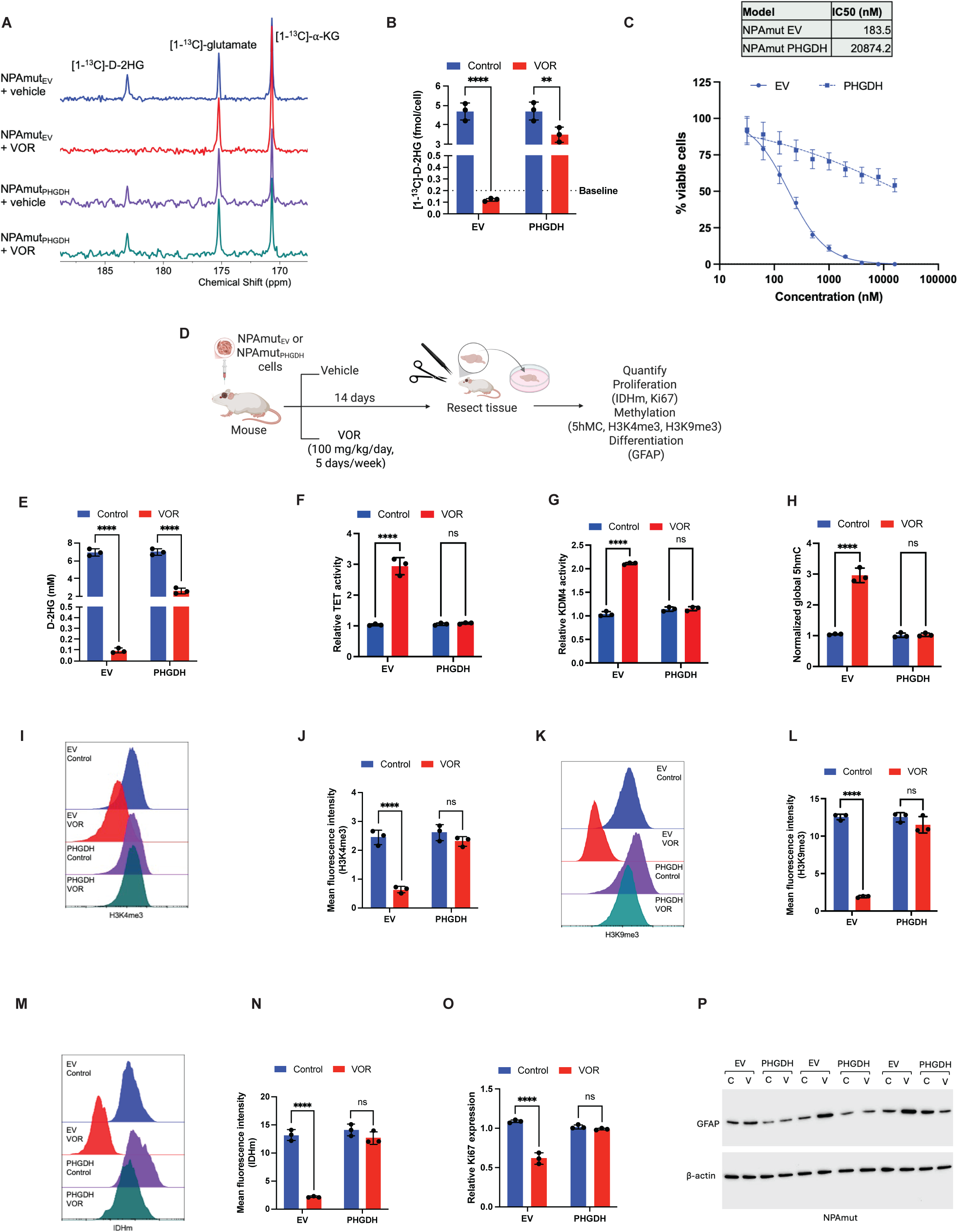
PHGDH drives D-2HG synthesis from α-KG in genetically engineered NPAmut cells. Representative ^13^C-MR spectra of [1-^13^C]-α-KG metabolism **(A)** and quantification of ^13^C-labeled D-2HG synthesis **(B)** in NPAmut cells overexpressing an EV (NPAmut_EV_) or PHGDH (NPAmut_PHGDH_). Cells were treated with vehicle or VOR for 72 h. **(C)** Dose response curves for VOR in NPAmut_EV_ and NPAmut_PHGDH_ cells. **(D)** Schematic illustration of study design with NPAmut tumors. Mice were intracranially implanted with NPAmut_EV_ or NPAmut_PHGDH_ cells. Once tumors were MR-detectable, mice were treated with VOR for 14 days and tumor tissue resected for *ex vivo* assays. D-2HG concentration **(E)**, TET DNA 5mC hydroxylase activity **(F)**, and KDM4 demethylase activity **(G)**, and global 5hmC abundance **(H)** in NPAmut_EV_ and NPAmut_PHGDH_ tumors treated with vehicle or VOR as described in panel D. Representative histograms **(I)** and quantification **(J)** of histone H3K4me3 in NPAmut_EV_ or NPAmut_PHGDH_ tumors treated with vehicle or VOR as described in panel D. Representative histograms **(K)** and quantification **(L)** of H3K9me3 expression in NPAmut_EV_ or NPAmut_PHGDH_ tumors treated with vehicle or VOR as described in panel D. Representative histograms **(M)** and quantification **(N)** of IDHm expression in NPAmut_EV_ or NPAmut_PHGDH_ tumors treated with vehicle or VOR as described in panel D. **(O)** Ki67 expression in NPAmut_EV_ or NPAmut_PHGDH_ tumors treated with vehicle or VOR as described in panel D. **(P)** Westerns blots for GFAP expression in NPAmut_EV_ or NPAmut_PHGDH_ tumors treated with vehicle or VOR as described in panel D. All studies were n≥3. Data is presented as mean ± standard deviation. * indicates statistical significance with p<0.05, ** indicates p<0.01, *** indicates p<0.001, **** indicates p<0.0001, ns is not significant.

To assess whether PHGDH mediates VOR resistance *in vivo,* we examined mice bearing intracranial NPAmut_EV_ or NPAmut_PHGDH_ tumors. Following treatment with vehicle or VOR for 14 days, tumor-bearing mice were euthanized, and tumor tissue was used for *ex vivo* analysis of pharmacodynamic biomarkers of response to VOR (see schematic in Fig. 4D). Consistent with our *in vitro* data, we observed a significant reduction in D-2HG abundance in VOR-treated NPAmut_EV_ tumors (Fig. 4E). In contrast, NPAmut_PHGDH_ tumors retained significant D-2HG abundance, despite treatment with VOR (Fig. 4E). Importantly, VOR caused a significant increase in TET and KDM4 enzyme activity, DNA hydroxylation, and a significant reduction in the proportion of cells with histone hypermethylation in NPAmut_EV_ tumors but not VOR-treated NPAmut_PHGDH_ tumors (Fig. 4F-4L). Furthermore, these molecular changes were accompanied by a significant decrease in tumor proliferation as measured by IDHm and Ki67 expression and a reciprocal increase in astrocytic differentiation as measured by GFAP expression in VOR-treated NPAmut_EV_ but not NPAmut_PHGDH_ tumors (Fig. 4M-4P). These results confirm a role for PHGDH in driving D-2HG production from α-KG and inducing VOR resistance in an isogenic setting.

### Elevated PHGDH is associated with resistance to IDHm inhibition in IDHm glioma patients

To confirm the clinical relevance of our findings, we examined tumor tissue from IDHm glioma patients (3 oligodendroglioma and 4 astrocytoma; grades 2-4) who were treated with either VOR or IVO at UCSF (see Fig. 5A and Supplementary Table S1 for details of the study design and patient cohort). The phase 3 indigo trial established the time to next intervention as a clinically significant variable, with 85.6% patients receiving a placebo requiring the next treatment intervention by 18 months, as opposed to 47.4% of VOR-treated patients (19). We, therefore, defined patients who progressed within 18 months (patients SF12239, SF128319, and SF14058) as intrinsically resistant to VOR. In contrast, patients SF12098, SF12264, and SF13345 were defined as intrinsically sensitive to VOR since they underwent disease progression after >40 months. As shown in the representative MRI in Fig. 5B, we observed tumor progression within 7 months of treatment with IVO in patient SF12239 (VOR-resistant), while patient SF12264 (VOR-sensitive) remained stable up to 59 months on VOR.

**Figure 5.**
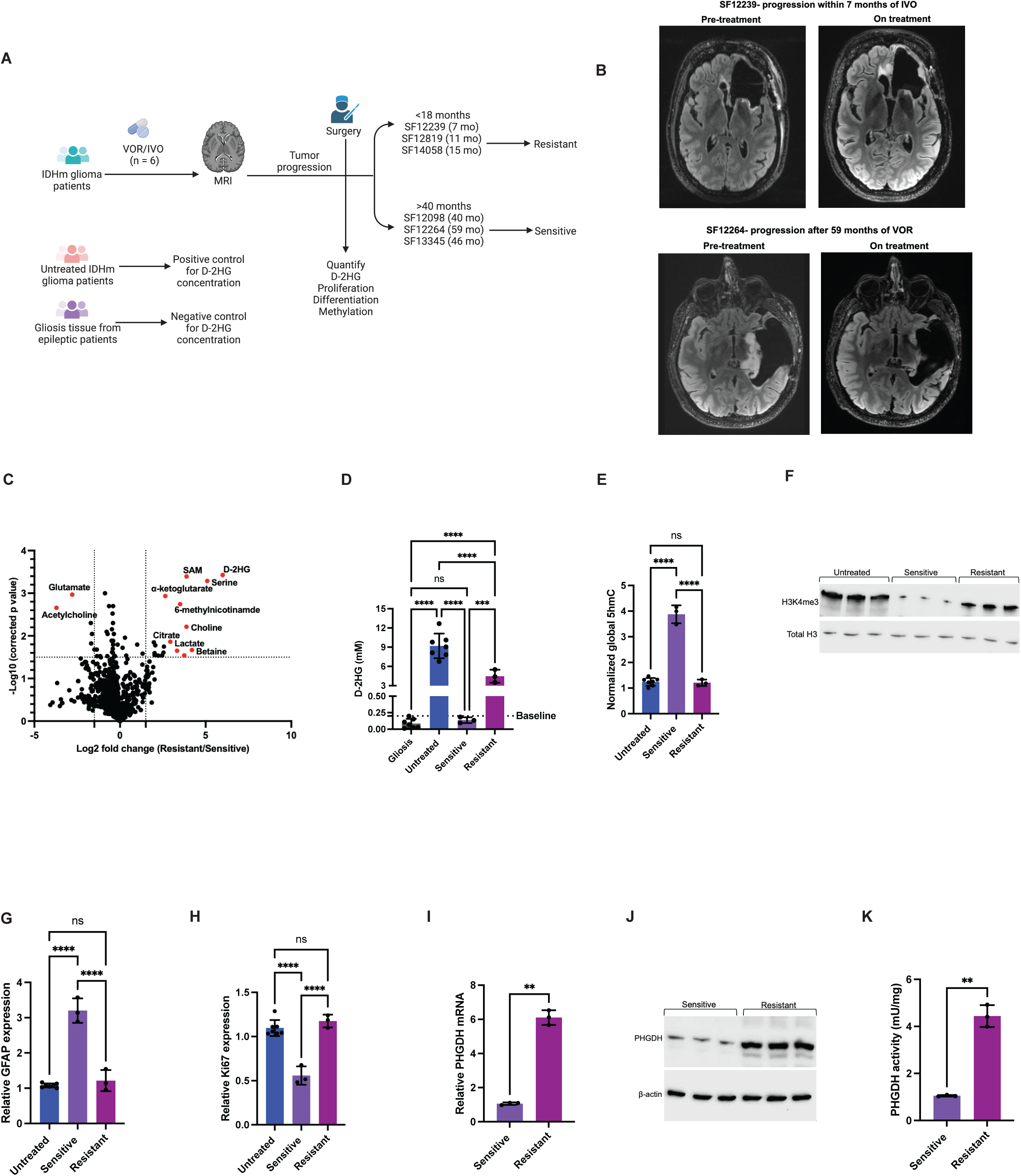
PHGDH is associated with intrinsic resistance to VOR in IDHm glioma patients. **(A)** Study design with VOR/IVO-sensitive and -resistant IDHm glioma patients. Tumor tissue resected from 3 patients who experienced disease progression within 18 months (resistant) and 3 patients whose tumors progressed after 40 months (sensitive) of treatment with VOR or IVO was used to measure D-2HG, proliferation, astrocytic differentiation, and methylation. Tumor tissue from patients who were never treated with VOR was used as external positive controls for D-2HG concentration, and gliosis tissue from epileptic patients was used as external negative controls. **(B)** *Top:* Axial gadolinium-enhanced T1-weighted MRI showing radiographic tumor progression in patient SF12239. *Bottom:* Axial gadolinium-enhanced T1-weighted MRI showing radiographic tumor regression in patient SF12264. **(C)** Volcano plot of differential metabolites in VOR/IVO-sensitive and VOR/IVO-resistant IDHm glioma patient tissue. **(D)** D-2HG concentration in VOR/IVO-sensitive and VOR/IVO-resistant tumor tissue. Untreated IDHm glioma and gliosis tissue were used as controls. **(E)** Global 5hmC abundance in VOR/IVO-sensitive, VOR/IVO-resistant, and gliosis tissue. **(F)** Western blots for histone H3K4me3 in VOR/IVO-sensitive, VOR/IVO-resistant, and untreated IDHm glioma patient tissue. mRNA expression for GFAP **(G)** and Ki67 **(H)** in VOR/IVO-sensitive, VOR/IVO-resistant, and untreated IDHm glioma patient tissue. PHGDH mRNA **(I)**, PHGDH protein **(J)**, and PHGDH activity **(K)** in VOR/IVO-sensitive and VOR/IVO-resistant IDHm glioma patient tissue. All studies were n≥3. Data is presented as mean ± standard deviation. * indicates statistical significance with p<0.05, ** indicates p<0.01, *** indicates p<0.001, **** indicates p<0.0001, ns is not significant.

First, we examined the global metabolomic profile of tumor tissue from VOR-sensitive and VOR-resistant patients using liquid chromatography mass spectrometry (LC-MS). As shown in Fig. 5C, VOR-resistant tumors had notably higher levels of D-2HG, serine, S-adenosylmethionine, 6-methylnicotinamide, lactate, choline, betaine, citrate, and lower levels of glutamate and acetylcholine relative to VOR-sensitive tumors. To further confirm a role for D-2HG, we compared D-2HG concentration in our patient cohort with external positive and negative controls. Due to the relative rarity of the patient population and the associated difficulties of obtaining matched pre- and post-treatment tumor tissue from the IVO/VOR-treated patients, we examined tumor tissue from grade-matched patients who were never treated with an IDHm inhibitor as external positive controls for D- 2HG abundance (see Supplementary Table S2) (15). In addition, we examined gliosis tissue from epileptic patients as external negative controls for D-2HG abundance. As shown in Fig. 5D, while treatment with VOR or IVO suppressed D-2HG abundance to baseline (defined here as the concentration observed in gliosis tissue, i.e., <0.2 mM) in VOR-sensitive patients, we observed significant D-2HG concentration (>2 mM) relative to baseline in tumor tissue from VOR-resistant patients. Concomitantly, global DNA hydroxylation was significantly elevated and histone H3K4me3 was significantly reduced in tumor tissue from VOR-sensitive but not VOR-resistant patients relative to untreated controls (Fig. 5E-5F). Functionally, we observed a significant increase in astrocytic differentiation and a reciprocal decrease in Ki67 expression in VOR-sensitive but not VOR-resistant patients relative to untreated controls (Fig. 5G-5H). Importantly, PHGDH mRNA, protein, and activity were significantly elevated in tumor tissue from VOR-resistant patients relative to VOR-sensitive patients (Fig. 5I-5K). Collectively, these results confirm the clinical relevance of PHGDH in mediating intrinsic resistance to IDHm inhibition in IDHm glioma patients.

### DMI of diethyl-[3,3’-^2^H]-α-KG metabolism to D-2HG provides a readout of IDHm expression in isogenic and patient-derived glioma cells

Imaging IDHm gliomas and their response to therapy, or lack thereof, is a significant challenge (20,23–25). Given that D-2HG accumulates to millimolar concentrations in IDHm glioma cells and is virtually undetectable in the normal brain, we questioned whether tracing the metabolism of diethyl-[3,3’-^2^H]-α-KG using DMI has the potential to provide a readout of IDHm activity in glioma cells. The decision to examine the diethyl ester of α-KG is dictated by the fact that α-KG is a dianion at physiological pH and cannot readily cross membranes. It is also unlikely to be readily transported across the BBB or the cell membrane by monocarboxylate transporters (46). Esterification is a widely used strategy to facilitate the membrane permeability of charged molecules, including α-KG (46–48). Following diffusion across the BBB and the cell membrane, the diethyl group is cleaved by ubiquitous cellular esterases to release [3,3′-^2^H]-α-KG within the cell. The choice of labeling at the 3 and 3’ positions for α-KG is based on the fact that it is expected to produce a single peak for α-KG at 3 ppm, thereby reducing the complexity of ^2^H-MR spectra, and allowing sufficient separation from the product (D-2HG; 1.9 ppm). Importantly, α-KG labeling at the 3 and 3’ positions provides a strategy for precision imaging of D-2HG production in IDHm gliomas (see schematic in Fig. 6A). Specifically, since α-KG conversion to D-2HG (1.9 ppm) is not reversible under physiological conditions (49,50), we expect accumulation of ^2^H-D-2HG in the tumor in IDHm gliomas. In contrast, [3,3′-^2^H]-α-KG metabolism via the tricarboxylic acid (TCA) cycle in the normal brain or in IDHwt tumors is expected to lead to loss of the ^2^H label during the conversion of succinate to fumarate by fumarase and during the conversion of isocitrate to α-KG by the IDHwt enzyme (Fig. 6A). Alternately, [3,3′-^2^H]-α-KG can be converted to ^2^H-glutamate (denoted by convention as glx; 2.4 ppm) by transamination in both IDHm and IDHwt tumors and normal brain.

**Figure 6.**
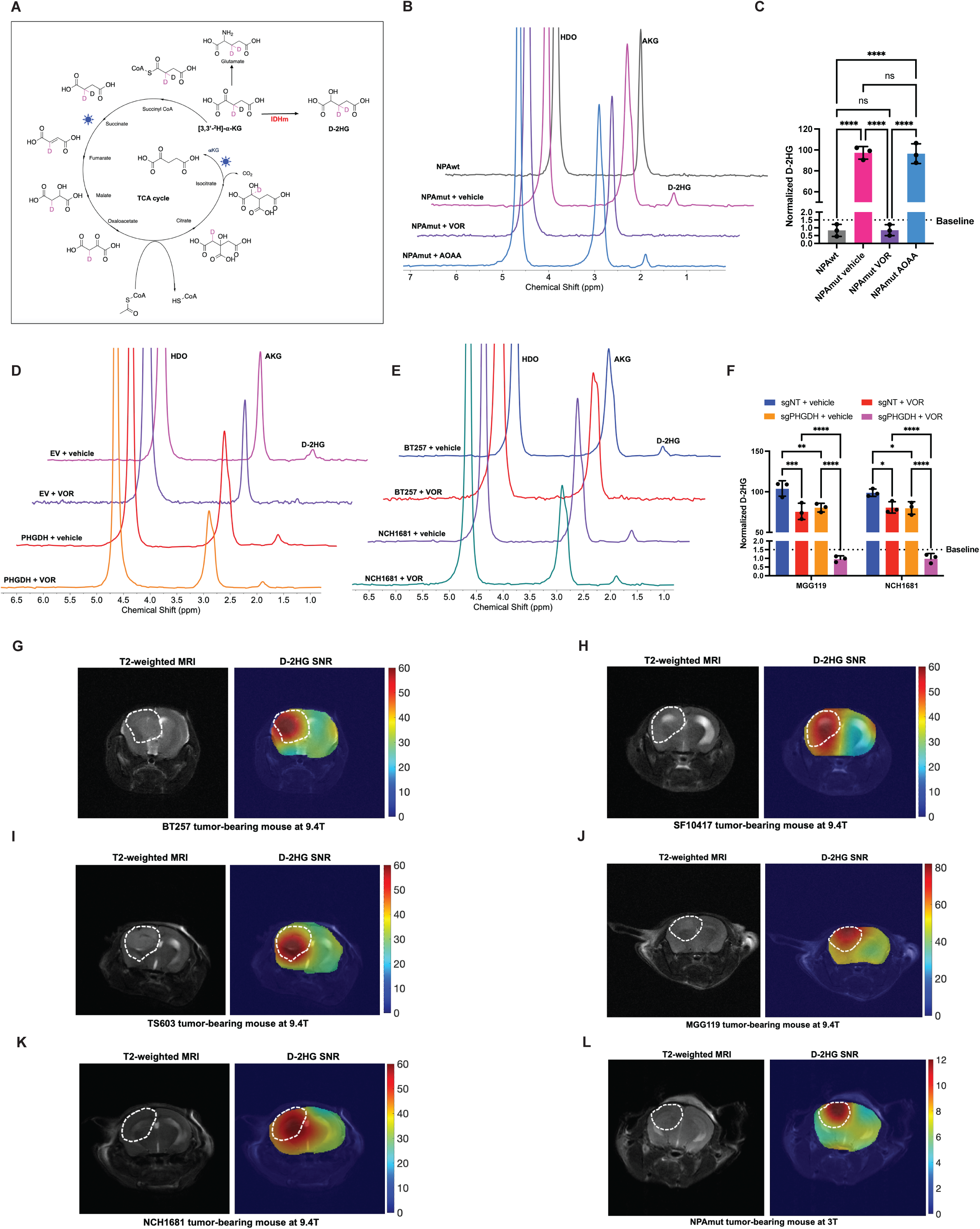
Diethyl-[3,3’-^2^H]-α-KG provides a readout of tumor burden in IDHm gliomas. **(A)** Schematic illustration of the rationale for using diethyl-[3,3’-^2^H]-α-KG in IDHm gliomas. * represent the enzymatic reactions where deuterons are lost. Representative ^2^H-MR spectra **(B)** and quantification **(C)** of diethyl-[3,3’-^2^H]-α-KG metabolism to D-2HG in NPAwt or NPAmut cells. NPAmut cells were treated with VOR or the transaminase inhibitor aminooxyacetate (AOAA) to assess whether the peak at 1.9 ppm corresponds to D-2HG or glx. **(D)** Representative ^2^H-MR spectra of diethyl-[3,3’-^2^H]-α-KG metabolism to D-2HG in NPAmut_EV_ and NPAmut_PHGDH_ cells treated with vehicle or VOR. **(E)** Representative ^2^H-MR spectra of diethyl-[3,3’-^2^H]-α-KG metabolism in patient-derived VOR-sensitive (BT257) or VOR-resistant (NCH1681) cells treated with vehicle or VOR. **(F)** Quantification of D-2HG production from diethyl-[3,3’-^2^H]-α-KG in MGG119 or NCH1681 cells expressing a non-targeting sgRNA or PHGDH-targeted sgRNA and treated with vehicle or VOR. **(G-K)** Representative 2D CSI data acquired at 9.4T in a mouse bearing an intracranial BT257 **(G)**, SF10417 **(H)**, TS603 **(I),** MGG119 **(J)**, or NCH1681 **(K)** tumor. In each case, the left panel shows the T2-weighted MRI with the tumor outlined with white dotted lines, and the right panel shows the heatmap of the SNR of D-2HG. **(L)** Representative 2D CSI data acquired at 3T in a mouse bearing an intracranial NPAmut tumor. The left panel shows the T2-weighted MRI with the tumor outlined with white dotted lines, and the right panel shows the heatmap of the SNR of D-2HG. All studies were n≥3. Data is presented as mean ± standard deviation. * indicates statistical significance with p<0.05, ** indicates p<0.01, *** indicates p<0.001, **** indicates p<0.0001, ns is not significant.

First, we compared the metabolism of non-esterified [3,3’-^2^H]-α-KG with diethyl-[3,3’-^2^H]-α-KG in NPAmut cells. As shown in Supplementary Fig. S3A-S3B, D-2HG production could be observed in NPAmut cells incubated with diethyl-[3,3’-^2^H]-α-KG but not with [3,3’-^2^H]-α-KG. We then examined diethyl-[3,3’-^2^H]-α-KG metabolism in NPAmut cells and compared it to isogenic cells expressing the IDHwt enzyme (NPAwt). As shown in the representative spectra in Fig. 6B and the quantification in Fig. 6C, while we observed a peak for α-KG at 3 ppm in both NPAwt and NPAmut cells, we saw a peak for D-2HG (1.9 ppm) specifically in NPAmut but not NPAwt cells. Importantly, treatment with VOR reduced the peak at 1.9 ppm to baseline levels observed in NPAwt cells, confirming that it corresponds to D-2HG (Fig. 6B-6C). Of note, we did not observe a peak for ^2^H-glx (2.4 ppm) in NPAwt or NPAmut cells (Fig. 6B-6C). To further confirm that the peak at 1.9 ppm corresponds to ^2^H-D-2HG and not ^2^H-glx, we treated NPAmut cells with aminooxy acetate, which is a pan-transaminase inhibitor. As shown in Fig. 6B-6C, treatment with aminooxy acetate did not impact the peak at 1.9 ppm in NPAmut cells. Collectively, these results indicate that metabolism of diethyl-[3,3’-^2^H]-α-KG to ^2^H-D-2HG is a specific readout of IDHm status in isogenic glioma cells.

### Diethyl-[3,3’-^2^H]-α-KG provides a readout of intrinsic resistance to IDHm inhibition in patient-derived glioma cells

Given our findings that PHGDH-mediated restoration of D-2HG production drives intrinsic resistance to IDHm inhibition, we questioned whether diethyl-[3,3’-^2^H]-α-KG provides a readout of intrinsic resistance to IDHm inhibitors in our glioma models. As shown in Fig. 6D and Supplementary Fig. S3C, while VOR abrogated ^2^H-D-2HG production in NPAmut_EV_ cells, NPAmut_PHGDH_ cells produced ^2^H-D-2HG despite treatment with VOR. Importantly, treatment with VOR inhibited diethyl-[3,3’-^2^H]-α-KG metabolism to ^2^H-D-2HG in patient-derived IDHm glioma cells that were intrinsically sensitive to VOR (BT257) but not those that were resistant to VOR (NCH1681; Fig. 6E and Supplementary Fig. S3D). Notably, silencing PHGDH blocked ^2^H-D-2HG production in VOR-resistant MGG119 and NCH1681 cells (Fig. 6F). These results are consistent with our mechanistic data and suggest that diethyl-[3,3’-^2^H]-α-KG has the potential to non-invasively interrogate intrinsic resistance to VOR in IDHm glioma cells.

### Spatially mapping D-2HG production from diethyl-[3,3’-^2^H]-α-KG enables visualization of the metabolically active tumor in mice bearing intracranial IDHm xenografts *in vivo*

Visualizing the spatial distribution of tumor burden is challenging in IDHm gliomas (20). We questioned whether quantifying the spatial distribution of ^2^H-D-2HG by overlaying the SNR of ^2^H-D-2HG over the corresponding T2-weighted MRI using 2D chemical shift imaging (CSI) would enable visualization of metabolically active tumor tissue in IDHm gliomas *in vivo*. As shown in Fig. 6G-6K and Supplementary Fig. S3E, while tumors were difficult to visualize by MRI, the metabolic heatmap of ^2^H-D-2HG produced from diethyl-[3,3’-^2^H]-α-KG clearly demarcated the tumor from the surrounding normal brain in mice bearing intracranial BT257, SF10417, TS603, MGG119, or NCH1681 tumors at a magnetic field strength of 9.4T. Importantly, ^2^H-D-2HG discriminated tumor from the contralateral normal brain in mice bearing intracranial NPAmut tumors at the clinically relevant magnetic field strength of 3T (Fig. 6L and Supplementary Fig. S3F). These results identify diethyl-[3,3’-^2^H]-α-KG as a unique tracer of ^2^H-D-2HG production in preclinical IDHm glioma models *in vivo*.

### Diethyl-[3,3’-^2^H]-α-KG has the potential to stratify VOR-sensitive and VOR-resistant IDHm gliomas *in vivo*

Accurately quantifying response to therapy is challenging in patients with IDHm gliomas, especially following treatment with cytostatic drugs such as VOR (23–25). To assess whether diethyl-[3,3’-^2^H]-α-KG has the potential to assess response to therapy in IDHm gliomas *in vivo,* we treated mice bearing intracranial TS603 (VOR-sensitive) or MGG119 (VOR-resistant) tumors with vehicle or VOR, and quantified diethyl-[3,3’-^2^H]-α-KG metabolism at day 5±2 (TS603) or day 4±3 (MGG119). Mice were then euthanized, and tumor tissue was resected for *ex vivo* assays (see schematic in Fig. 7A). ^2^H-D-2HG production from diethyl-[3,3’-^2^H]-α-KG was significantly reduced in VOR-treated TS603 mice relative to vehicle controls at day 5±2 (Fig. 7B-7C), when no change in tumor volume could be detected by MRI (Supplementary Fig. S4A). Importantly, this drop in ^2^H-D-2HG production was associated with a significant reduction in the proportion of IDHm+ cells and a reciprocal increase in GFAP+ cells in VOR-treated TS603 tumors relative to controls (Fig. 7D-7E). In contrast, we observed a significant, albeit modest, increase in ^2^H-D-2HG production in VOR-treated MGG119 tumors relative to vehicle controls at day 4±3 (Fig. 7F-7G). As with TS603 tumors, there was no significant change in tumor volume at this early time point between VOR-treated and vehicle-treated MGG119 tumors (Supplementary Fig. S4B). Furthermore, we observed a significant increase in the proportion of IDHm+ cells and a reciprocal decrease in GFAP+ cells in VOR-treated MGG119 tumors relative to controls (Fig. 7H-7I).

**Figure 7.**
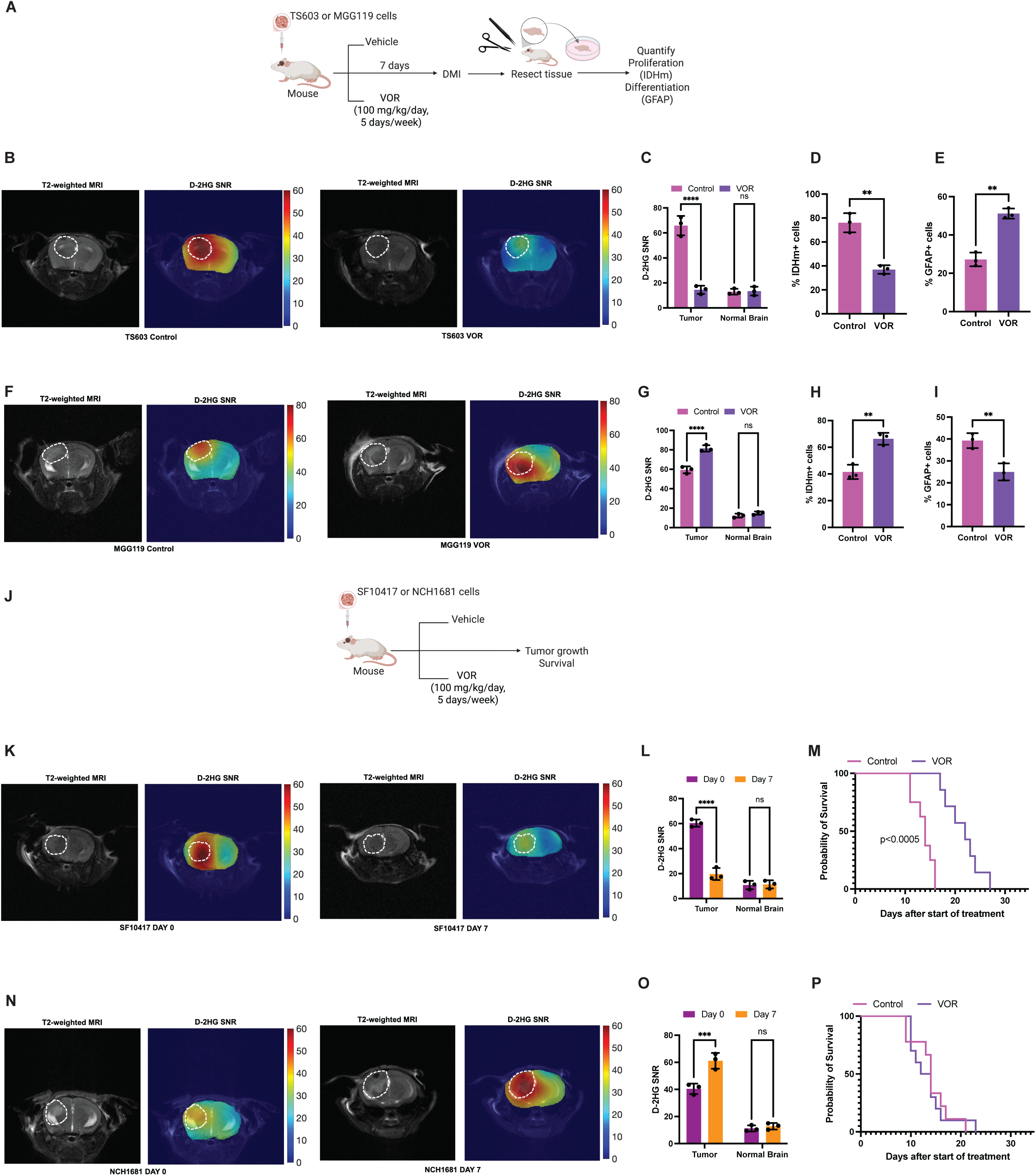
Diethyl-[3,3’-^2^H]-α-KG reports on early response to therapy and intrinsic resistance to VOR in IDHm gliomas. **(A)** Schematic illustration of the study designed to assess whether diethyl-[3,3’-^2^H]-α-KG provides a pharmacodynamic readout of treatment response in IDHm glioma-bearing mice. Mice bearing intracranial VOR-sensitive (TS603) or VOR-resistant (MGG119) tumors were treated with vehicle or VOR for ∼7 days. Following the acquisition of 2D CSI data, mice were euthanized and tumor tissue resected for *ex vivo* assays. **(B)** Representative 2D CSI data acquired at day 4 from mice bearing intracranial TS603 tumors treated with vehicle or VOR as described in panel A. In each case, the left panel shows the T2-weighted MRI with the tumor outlined in white dotted lines, and the right panel shows the heatmap of the SNR of ^2^H-D-2HG. Quantification of the SNR of ^2^H-D-2HG **(C)**, the % of IDHm+ cells **(D)**, and the % of GFAP+ cells **(E)** in TS603 mice treated with vehicle or VOR as described in panel A. **(F)** Representative 2D CSI data acquired at day 5 from mice bearing intracranial MGG119 tumors treated with vehicle or VOR as described in panel A. In each case, the left panel shows the T2-weighted MRI with the tumor outlined in white dotted lines, and the right panel shows the heatmap of the SNR of ^2^H-D-2HG. Quantification of the SNR of ^2^H-D-2HG **(G)**, the % of IDHm+ cells **(H)**, and the % of GFAP+ cells **(I)** in MGG119 mice treated with vehicle or VOR as described in panel A. **(J)** Schematic illustration of the study designed to assess whether diethyl-[3,3’-^2^H]-α-KG enables early stratification of responders from non-responders *in vivo.* Mice bearing intracranial VOR-sensitive (SF10417) or VOR-resistant (NCH1681) tumors were treated with vehicle or VOR for 7 days. Following the acquisition of 2D CSI data, mice were treated until they needed to be euthanized and monitored for survival. **(K)** Representative 2D CSI data acquired before (day 0) and after (day 7) treatment of mice bearing intracranial SF10417 tumors with VOR as described in panel J. In each case, the left panel shows the T2-weighted MRI with the tumor outlined in white dotted lines, and the right panel shows the heatmap of the SNR of ^2^H-D-2HG. Quantification of the SNR of ^2^H-D-2HG **(L)**, and survival **(M)** of mice bearing intracranial SF10417 tumors treated with VOR as described in panel J. **(N)** Representative 2D CSI data acquired at day 7 from mice bearing intracranial NCH1681 tumors treated with vehicle or VOR as described in panel J. In each case, the left panel shows the T2-weighted MRI with the tumor outlined in white dotted lines, and the right panel shows the heatmap of the SNR of ^2^H-D-2HG. Quantification of the SNR of ^2^H-D-2HG **(O)**, and survival **(P)** of mice bearing intracranial NCH1681 tumors treated with VOR as described in panel J. All studies were n≥3. Data is presented as mean ± standard deviation. * indicates statistical significance with p<0.05, ** indicates p<0.01, *** indicates p<0.001, **** indicates p<0.0001, ns is not significant.

To further validate these results, we treated mice bearing intracranial VOR-sensitive (SF10417) or VOR-resistant (NCH1681) tumors with vehicle or VOR and quantified diethyl-[3,3’-^2^H]-α-KG at day 7±1. Mice were treated until they needed to be euthanized according to animal care guidelines (see schematic in Fig. 7J). As shown in Fig. 7K-7L and Supplementary Fig. S4C, ^2^H-D-2HG production from diethyl-[3,3’-^2^H]-α-KG was significantly reduced at day 7±1 relative to day 0 in VOR-treated SF10417 mice, at a time when MRI-detectable tumor volume was unaltered. Importantly, this reduction in ^2^H-D-2HG production preceded the significant increase in median survival (22 days for VOR-treated SF10417 mice vs. 14 days for vehicle-treated mice; Fig. 7M). Conversely, for the VOR-resistant NCH1681 model, ^2^H-D-2HG production from diethyl-[3,3’-^2^H]-α-KG was significantly elevated at day 7±1 relative to day 0 in VOR-treated mice (Fig. 7N-7O). MRI-detectable tumor volume was unaltered at this early timepoint in NCH1681 mice (Supplementary Fig. S4D). Importantly, consistent with the lack of a drop in ^2^H-D-2HG production, there was no change in survival between VOR-treated mice and vehicle-treated controls for the NCH1681 model (13 days for VOR-treated NCH1681 mice vs. 14 days for vehicle-treated mice; Fig. 7P). Collectively, these results indicate that tracing D-2HG production from diethyl-[3,3’-^2^H]-α-KG provides an early readout of treatment response that predicts survival benefit and can be used to stratify responders from non-responders *in vivo*.

## DISCUSSION

IDHm gliomas are malignant primary brain tumors that afflict relatively younger patients and have long-lasting and life-altering effects on physical and cognitive abilities (3–5). VOR was recently approved as a molecularly targeted therapy for IDHm glioma patients (19). Understanding the molecular mechanisms governing response to VOR and identifying imaging biomarkers that can accurately stratify responders from non-responders will facilitate the effective deployment of VOR to IDHm glioma patients (51). In this study, we demonstrate that restoration of D-2HG synthesis contributes to intrinsic resistance to VOR in IDHm gliomas. We show that the metabolic enzyme PHGDH promiscuously converts α-KG to D-2HG in IDHm gliomas with intrinsic resistance to VOR. Importantly, tracing D-2HG production from diethyl-[3,3’-^2^H]-α-KG using DMI enables non-invasive imaging of tumor burden and stratification of responders from non-responders *in vivo*.

Intrinsic or primary resistance to IVO has been reported in IDHm-driven acute myeloid leukemia (52). However, mechanisms driving resistance to IDHm inhibition in gliomas remain unclear. We examined the effect of VOR on viability using a panel of 12 patient-derived IDHm glioma models. Of note, these models harbor a variety of secondary and tertiary genetic alterations, and we did not observe any genomic alteration that is associated with resistance to VOR across these models. Since IDHm is a metabolic enzyme and α-KG participates in the TCA cycle, which produces ATP, we rigorously quantified the effect of VOR on viability using multiple non-metabolic readouts such as clonogenic potential and Ki67 expression. Our findings indicate that our patient-derived models can be stratified as VOR-sensitive (50%; IC50 ∼90-300 nM) and VOR-resistant (50%; IC50 >5-20 μM). Since all of these models were established from IDHm glioma patients who had never been treated with an IDHm inhibitor, these results confirm the presence of intrinsic resistance to VOR in IDHm glioma patients. It is also important to note that our patient-derived cells were treated with VOR for 72 h, which makes it unlikely that the resistance we observe is secondary or acquired resistance. A caveat with our patient-derived models is that they were largely derived from patients with grade 3 or 4 tumors (although GSC580 was isolated from a patient with a grade 2 astrocytoma, it bears a CDKN2A/2B deletion, which classifies it as a “molecular” grade 4 tumor). In contrast, the INDIGO trial was restricted to patients with grade 2 gliomas (19). To address this limitation, we examined tumor tissue resected from IDHm glioma patients, including those who were originally diagnosed with grade 2 tumors. These patients were treated with VOR or IVO and subsequently experienced disease progression. Given that the INDIGO trial demonstrated that the majority of placebo-treated patients received the next intervention within 18 months (19), we stratified patients who received the next intervention in <18 months as VOR-resistant and those with the next intervention in >40 months as VOR-sensitive. Consistent with the data from our patient-derived models, we observed a significant reduction in Ki67 expression in VOR-sensitive but not VOR-resistant patient tissue relative to untreated external controls. Although it is likely that acquired resistance to VOR will emerge as more patients receive treatment with VOR, nonetheless, our studies demonstrate the existence of intrinsic resistance to IDHm inhibition in IDHm gliomas.

We identify, to the best of our knowledge, for the first time, PHGDH-driven restoration of D-2HG synthesis as a molecular driver of intrinsic resistance to IDHm inhibition in gliomas. Stable isotope tracing demonstrated that VOR reduced [1-^13^C]-D-2HG synthesis from [1-^13^C]-α-KG to baseline levels observed in primary human astrocytes in patient-derived IDHm glioma models that are intrinsically sensitive to VOR. In contrast, patient-derived models that are intrinsically resistant to VOR retained substantial [1-^13^C]-D-2HG synthesis from [1-^13^C]-α-KG, even in the presence of VOR, pointing to another source of D-2HG in these cells. Similarly, D-2HG concentration was reduced to baseline levels observed in gliosis tissue in tumor tissue from VOR-sensitive IDHm glioma patients. In contrast, D-2HG concentration was significantly elevated relative to baseline in tumor tissue from VOR-resistant IDHm glioma patients. Guided by previous studies showing that PHGDH promiscuously converts α-KG to D-2HG in bacterial and mammalian cells (40,41), we examined PHGDH expression in our models. PHGDH expression was significantly higher in VOR-resistant patient-derived models relative to VOR-sensitive models. Silencing PHGDH in combination with VOR reduced [1-^13^C]-D-2HG synthesis to baseline in VOR-resistant cells. Conversely, overexpressing PHGDH restored [1-^13^C]-D-2HG synthesis despite treatment with VOR in VOR-sensitive cells. Importantly, PHGDH expression was significantly higher in VOR-resistant patient tissue relative to tissue from VOR-sensitive patients. Furthermore, consistent with the normal physiological role of PHDGH in driving serine synthesis from glucose-derived 3-phosphoglycerate (53), untargeted metabolomics of patient tissue identified elevated serine and D-2HG in VOR-sensitive tumors relative to VOR-resistant tumors. Although further studies are needed to identify the molecular mechanisms driving elevated baseline PHGDH expression in IDHm gliomas, our studies nonetheless pinpoint a role for PHGDH in contributing to VOR resistance in IDHm gliomas.

Our studies identify PHGDH as a collateral therapeutic vulnerability in VOR-resistant gliomas. Overexpressing PHGDH induces VOR resistance in IDHm glioma models, while conversely, silencing PHGDH confers sensitivity to VOR in IDHm gliomas. It is important to note that while silencing PHGDH in VOR-resistant tumors reduces the IC50 value from 7-20 μM to 0.3-0.8 μM, the IC50 values do not always reach levels observed in VOR-sensitive models (<0.3 μM), pointing to the presence of additional mechanisms of resistance to VOR. Indeed, untargeted metabolomics of patient tissue identifies elevated levels of other metabolites such as S-adenosylmethionine and lactate in VOR-resistant patient tissue. Further studies are warranted to assess if these metabolites or their associated pathways contribute to VOR resistance in IDHm gliomas.

We show, for the first time, that diethyl-[3,3’-^2^H]-α-KG is a novel, non-invasive tracer of D-2HG synthesis in IDHm gliomas. Due to its key role in driving tumorigenesis and its unique presence in IDHm tumors, D-2HG provides an unparalleled opportunity to devise non-invasive methods of imaging IDHm gliomas. ^1^H-MRS quantifies metabolite concentration, as opposed to synthesis, and has been used to detect D-2HG in IDHm glioma patients and to assess response to therapy (26,54–57). However, unambiguous D-2HG detection by ^1^H-MRS is complicated by its complex spectral pattern and overlap with metabolites that resonate with a similar chemical shift, such as glutamate, glutamine, and γ-aminobutyric acid, and are abundant in the normal brain (58). Hyperpolarized ^13^C-MRS following administration of [1-^13^C]-α-KG has been used to quantify dynamic conversion of D-2HG in preclinical IDHm glioma models (59). However, in addition to logistical challenges relating to rapid substrate administration and the need for an expensive hyperpolarizer, transport across the BBB is a severe limitation for hyperpolarized [1-^13^C]-α-KG due to the inability of α-KG to cross the cell membrane and the need to detect metabolism within <5 minutes. Indeed, we established that esterification was essential for imaging since D-2HG production could only be detected from diethyl-[3,3’-^2^H]-α-KG and not from [3,3’-^2^H]-α-KG. Importantly, we robustly demonstrate that 2D CSI of D-2HG production from diethyl-[3,3’-^2^H]-α-KG provides a spatial map of D-2HG production in real time in mice bearing intracranial IDHm glioma xenografts from multiple patient-derived IDHm glioma models. Furthermore, tumors that were difficult to visualize by MRI were clearly demarcated from the surrounding normal brain by DMI with excellent tumor-to-background contrast *in vivo*. Collectively, our studies highlight the potential of diethyl-[3,3’-^2^H]-α-KG for visualizing tumor burden in IDHm gliomas and suggest that tracing D-2HG production from diethyl-[3,3’-^2^H]-α-KG can be used to non-invasively identify IDHm glioma patients for enrollment in window-of-opportunity clinical trials (60).

Following the onset of therapy, accurately assessing whether the patient is responding to treatment and stratifying responders from non-responders at an early time point remains a challenge with IDHm glioma patients (23–25). This is particularly true for precision therapies such as VOR that do not always induce tumor regression. Our studies indicate that D-2HG production from diethyl-[3,3’-^2^H]-α-KG is significantly reduced within a week of treatment with VOR in mice bearing intracranial IDHm gliomas that are intrinsically sensitive to VOR. In contrast, we observed a significant increase in D-2HG production from diethyl-[3,3’-^2^H]-α-KG after a week of treatment with VOR in mice bearing intracranial IDHm gliomas that are intrinsically resistant to VOR. With regard to clinical translation, our DMI studies were performed both at 9.4T and the widely available clinical field strength of 3T, which facilitates widespread dissemination. Taken together, our studies provide proof-of-concept evidence that DMI using diethyl-[3,3’-^2^H]-α-KG provides a readout of early response to therapy in IDHm gliomas and enables stratification of responders from non-responders.

In summary, using clinically relevant patient-derived IDHm glioma models and patient tissue, we demonstrate that PHGDH-driven D-2HG synthesis mediates intrinsic resistance to VOR in IDHm gliomas. Importantly, we show that diethyl-[3,3’-^2^H]-α-KG is a unique tool for the non-invasive assessment of tumor burden, treatment response, and patient stratification in IDHm gliomas.

## MATERIALS AND METHODS

### Cell culture

All IDHm glioma cells were maintained as neurospheres in serum-free Neurobasal-A medium (Gibco) supplemented with glutamine, growth factors, and antibiotics (29–36,42). BT257, BT142, and BT54 cells were a kind gift from Dr. Luchman (University of Calgary). SF10417 and SF10602 cells were a kind gift from Dr. Costello (UCSF). TS603 cells were kindly provided by Dr. Venneti (University of Michigan, Ann Arbor). MGG119 and MGG152 cells were a kind gift from Dr. Cahill (Massachusetts General Hospital). NCH1681 and NCH612 were kindly gifted by Dr. Herold-Mende (University of Heidelberg). GSC580 cells were a kind gift from Dr. Lamfers (Erasmus University). NPAmut and NPAwt cells were kindly provided by Dr. Castro (University of Michigan, Ann Arbor). GBM6 cells were obtained from the Mayo Clinic Biorepository. All cell lines were routinely tested for mycoplasma contamination, authenticated by short tandem repeat fingerprinting, and assayed within 6 months of authentication.

### IC50

Colony formation was determined as described previously (61). Neurospheres were disaggregated using Accutase. Single cells were mixed with collagen and plated into a 6-well plate at 7,500 cells per well in serum-free medium containing growth factors. Cells were treated with vehicle or a 10-point serial dilution of VOR. The wells were replenished with fresh medium containing VOR every 3 days. After 21 days, colonies were counted and the % inhibition of clonogenicity was measured. Ki67 expression was measured using the Lumit hKi67 assay (Promega) according to the manufacturer’s instructions.

### Activity assays

D-2HG concentration was measured using a commercially available kit that is highly specific for D-2HG (Abcam, ab211070) (37). Briefly, cell or tissue samples are homogenized and mixed with a reaction mix containing D2HGDH, which oxidizes D-2HG to α-KG. Concomitantly, NAD+ is reduced to NADH, which is then quantified by the diaphorase-mediated reduction of rezasurin to resorufin. TET 5mC hydroxylase and KDM4 DNA demethylase activity were similarly quantified using kits (Abcam, ab156913 and ab113462 respectively). Global 5hMC abundance was quantified using an ELISA-based kit (Active Motif, 55018). PHGDH activity was quantified using a kit (Abcam, ab273328).

### Stable isotope tracing in cells

5×10^5^ cells were seeded in 6-well plates and incubated in cell culture media containing 2 mM [1-13C]-α-KG for 72 h. Cell resuspensions were mixed with a known concentration of ^13^C-labeled methanol and proton-decoupled ^13^C-MR spectra were acquired (30° flip angle, TR = 3s, 2048 acquisitions) using a 14.1 T Bruker 600 spectrometer. Peak integrals were calculated using Mnova and normalized to the external reference (^13^C-methanol) and cell number.

### Gene silencing and overexpression

To silence PHGDH in IDHm glioma cells, we used Edit-R All-in-one lentiviral particles expressing a pool of 3 sgRNA targeting PHGDH and the Cas9 nuclease under the control of the hEF1a promoter and harboring a puromycin resistance marker. Lentiviral particles expressing a non-targeting control sequence (sgNT) were used as controls. IDHm glioma cells were transduced with the lentivirus in serum-free medium using polybrene and selected using puromycin. PHGDH knockdown was verified using quantitative PCR (QPCR) and immunoblotting. To overexpress PHGDH, IDHm glioma cells were transfected with lentiviral particles expressing Myc-tagged human PHGDH or the corresponding empty vector using Turbofectin and selected using puromycin. PHGDH expression was verified by QPCR.

### QPCR

Gene expression was measured by QPCR on a QuantStudio 5 system (ThermoFisher), and data was normalized to β-actin. Total RNA was extracted using the TRIzol reagent and converted to cDNA using the High-Capacity cDNA Reverse Transcription Kit (ThermoFisher Scientific). QPCR was performed using the TaqMan Fast Advanced Master Mix (ThermoFisher Scientific) and TaqMan assay primers.

### Patient biopsies

Patient biopsies were obtained from the UCSF Brain Tumor Center Biorepository in compliance with the written informed consent policy. Biopsy use was approved by the Committee on Human Research at UCSF, and research was approved by the Institutional Review Board at UCSF according to ethical guidelines established by the U.S. Common Rule.

### Metabolomics

Metabolites were extracted from ∼10 mg of tissue using 4:1 methanol: water. LC-MS was performed using an ultra-performance LC (ExionLC 2.0) and Quadrupole-Time of Flight Spectrometry (TripleTOF 6600+, AB SciEx). Metabolites were separated using a Waters ACQUITY T3 C18 column (1.8 μm, 2.1 mm x 100 mm) column with mobile phases A (0.1 % formic acid in water) and B (0.1% formic acid in acetonitrile). High-resolution MS scans were acquired in positive and negative modes. Metabolites were identified by comparison to standards. Peak areas were calculated, corrected to blank samples, and normalized to the protein content of the sample.

### Immunoblotting

Cells or tissue were lysed using Cell Lytic Buffer (Sigma) and separated on a 4-20% polyacrylamide gel (Bio-Rad). Following transfer to a PVDF membrane, proteins were detected by staining with the relevant primary antibody, followed by incubation with the corresponding HRP-conjugated secondary antibody. Proteins were visualized using an enhanced chemiluminescence substrate kit (ThermoFisher) and the Chemisolo imager (Azure Biosciences). Blots were probed for H3K4me3 (ThermoFisher, MA5-11199), H3K9me3 (ThermoFisher, 49-1008), PHGDH (Sigma, HPA021241), and GFAP (ThermoFisher, 13-0300). β-actin (Cell Signaling, 4970) was used as a loading control and goat anti-rabbit IgG-HRP (Cell Signaling, 7074) was used as a secondary antibody.

### Synthesis

Diethyl-[3,3’-^2^H]-α-KG was synthesized as described previously (48,62). Briefly, α-KG was deuterated to -[3,3’-^2^H]-α-KG using D2O at basic pH. Following the addition of concentrated sulfuric acid to a solution of -[3,3’-^2^H]-α-KG in ethyl alcohol, the reaction mixture was dried and concentrated to obtain diethyl-[3,3’-^2^H]-α-KG. ^2^H incorporation was quantified by mass spectrometry and by ^1^H-MRS.

### DMI of cell suspensions

2×10^6^ cells were incubated in growth media containing 2 mM diethyl-[3,3’-^2^H] α-KG for 72 h. For drug studies, cells were concurrently treated with vehicle (DMSO) or 1 μM VOR for 72 h. Cells were harvested, counted, and resuspended in growth media. ^2^H-MR spectra were acquired on a 14.1T spectrometer. The peak integral for D-2HG (1.9 ppm) was normalized to α-KG (3 ppm), HDO (4.75 ppm), and cell number.

### Tumors

All studies were approved by the University of California, San Francisco Institutional Animal Care and Use Committee (IACUC). Cells were intracranially injected into the cortex of 5-6 weeks old female C57BL/6 mice (NPAmut_EV_, NPAmut_PHGDH_) or SCID mice (BT257, SF10417, TS603, MGG119, NCH1681) (28,63). Intracranial implantation was performed using a stereotactic frame at 2 mm to the right of the medial suture, 2 mm behind the bregma, and a depth of 2 mm. Tumor volume was determined by T2-weighted MRI using a Bruker 3T or 9.4T scanner (63). Axial T2-weighted images were acquired using a spin-echo TurboRARE sequence (TE/TR = 8.25/3200 ms, FOV = 30 x 30 mm^2^, 256 x 256, slice thickness = 1.2 mm, NA = 3). Once tumors were detectable by MRI, mice were treated with vehicle (Oraplus) or VOR (100 mg/kg/day, 5 days/week in Oraplus) via oral gavage.

### DMI in vivo

Studies were performed on a Bruker 3T or 9.4T scanner using a 16 mm ^2^H surface coil. Following intravenous injection of a bolus of 2 g/kg of diethyl-[3,3’-^2^H]-α-ketoglutarate, 2D CSI data was acquired using the following parameters (TE/TR = 1.104/591.824 ms, FOV = 30 x 30 x 5 mm^3^, 512 points, spectral width = 925.9 Hz, NA = 8, temporal resolution = 5 minutes 3 s, nominal voxel size = 70.3 μL) at 9.4T. For 2D CSI at 3T, the following parameters were used (TE/TR = 1.04/265.89 ms, FOV = 30 x 30 x 8 mm^3^, complex points=128 points, spectral width = 2.5 kHz, NA = 30, temporal resolution = 8 minutes 30 s, nominal voxel size = 112.5 μL). Data was analyzed using MATLAB as described previously (28,63).

### Statistical analysis

All experiments were performed on a minimum of 3 biological replicates (n≥3), and the results were expressed as mean ± standard deviation. Statistical significance was assessed in GraphPad Prism 10 using a one-way ANOVA, two-way ANOVA, or two-tailed Welch’s t-test, with p<0.05 considered significant. Analyses were corrected for multiple comparisons using Tukey’s or Šídák’s multiple comparisons test, wherever applicable. Survival was quantified using Kaplan-Meier analysis. * indicates statistical significance with p<0.05, ** indicates p<0.01, *** indicates p<0.001 and **** indicates p<0.0001.

## Data availability

The data generated in this study are available from the corresponding author upon request.

## Acknowledgments

We thank Dr. Artee Luchman (University of Calgary), Dr. Joseph Costello (University of California, San Francisco), Dr. Dan Cahill (Massachusetts General Hospital), Dr. Sriram Venneti (University of Michigan), Dr. Christel Herold-Mende (University of Heidelberg), Dr. Martin Lamfers (Erasmus University), and Dr. Maria Castro (University of Michigan) for kindly providing the patient-derived models used in this study.

## Author Contributions

*Conceptualization, supervision, project administration, funding acquisition:* PV. *Development of methodology:* PV, CT, GB. *Investigation:* PV, CT, GB, AMG. *Formal Analysis:* PV, CT. *Software:* GB. *Resources:* JJP, JWT. *Writing – original draft:* PV. *Writing – review:* PV, CT, GB, JJP, JWT.

