## Supplementary Figures S1-S4 and Supplementary Table S1 for "Understanding and imaging PHGDH-driven intrinsic resistance to mutant IDH inhibition in gliomas"

Supplementary Figure 1

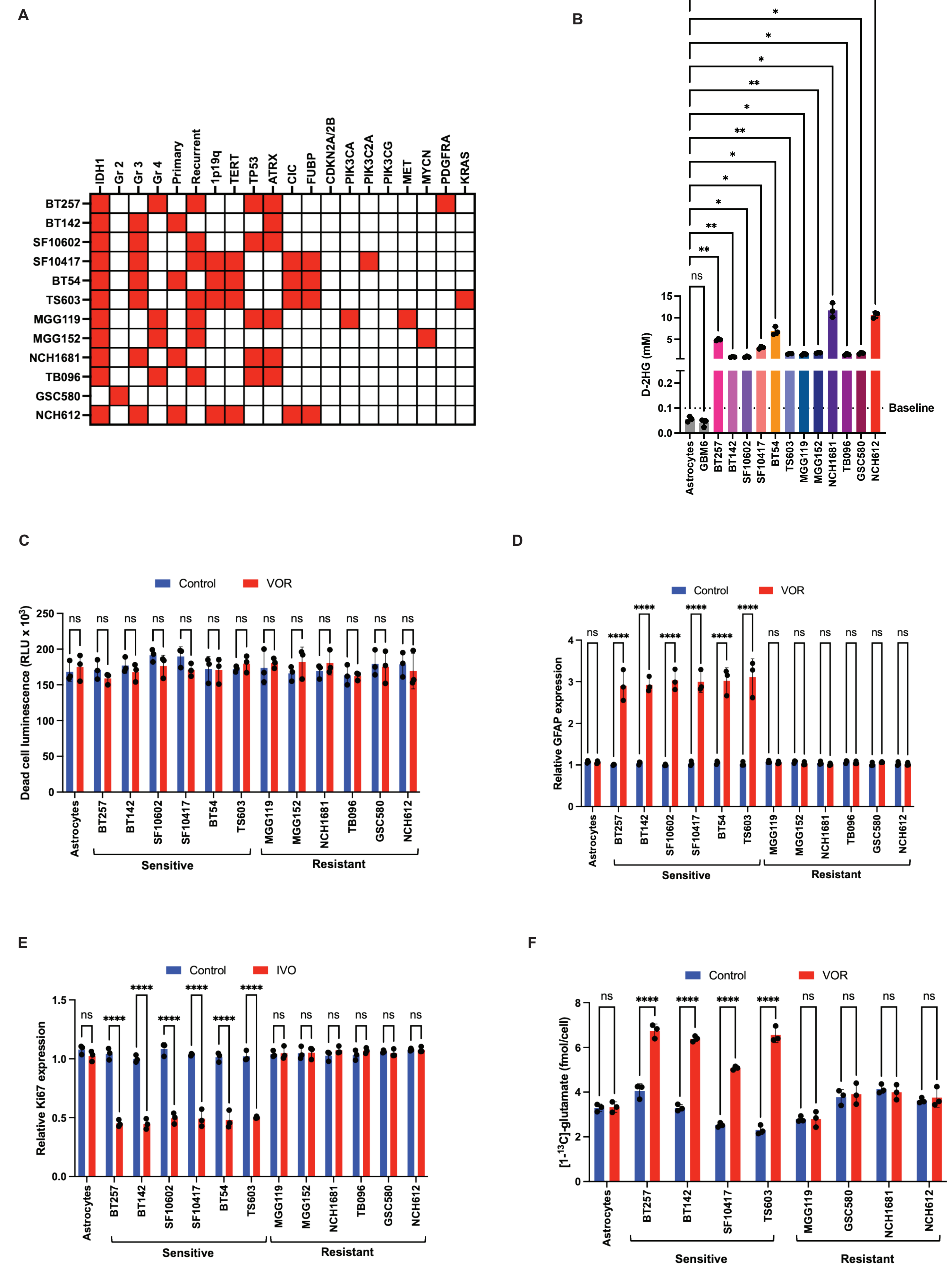

**Supplementary Figure 1. Intrinsic resistance to VOR occurs in patient-derived IDHm glioma models. (A)** Schematic representation of genetic and histological information available on the 12 patient-derived IDHm glioma models. Filled red squares indicate the presence of a genomic alteration. **(B)** D-2HG concentration in the patient-derived models used in this study. Primary human astrocytes were used as baseline controls. Effect of VOR on cytotoxicity **(C)** and astrocytic differentiation as measured by GFAP expression **(D)** in patient-derived IDHm glioma models. **(E)** Effect of IVO on viability as measured by Ki67 expression in patient-derived IDHm glioma models. **(F)** Quantification of [1-$^{13}$C]-glutamate production from [1-$^{13}$C]-$\alpha$-KG in VOR-sensitive (BT257, BT142, SF10417, TS603) and VOR-resistant (MGG119, GSC580, NCH1681, and NCH612) IDHm glioma cells. Primary human astrocytes were used as baseline controls. All studies were $n \geq 3$. Data is presented as mean ± standard deviation. * indicates statistical significance with $p<0.05$, ** indicates $p<0.01$, *** indicates $p<0.001$, **** indicates $p<0.0001$, ns is not significant.

### Supplementary Figure 2

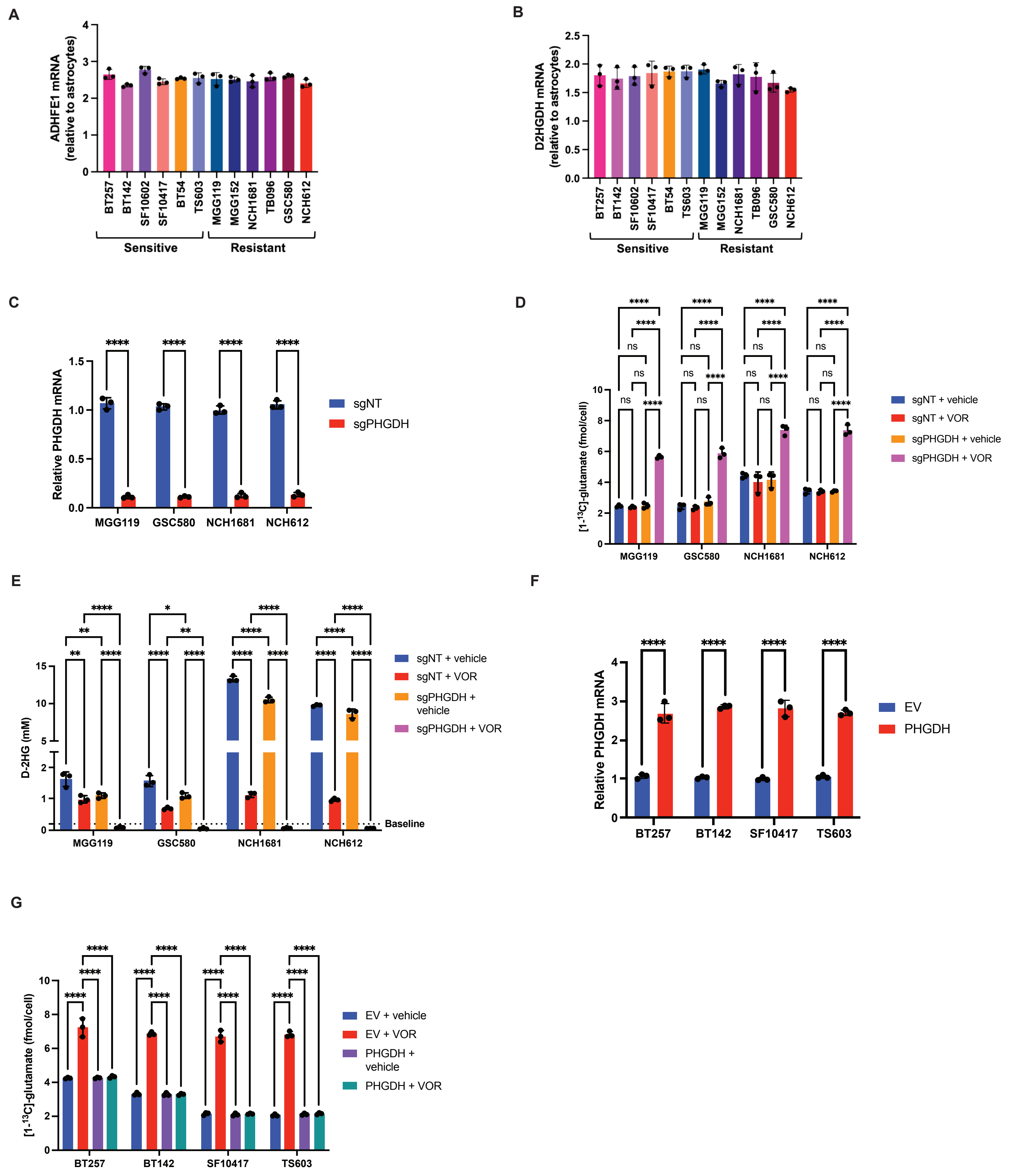

**Supplementary Figure 2. PHGDH contributes to intrinsic resistance to VOR in IDHm gliomas.** Quantification of ADHFE1 **(A)** and D2HGDH **(B)** mRNA in patient-derived IDHm glioma cells. **(C)** Verification of PHGDH silencing in VOR-resistant (MGG119, GSC580, NCH1681, NCH612) IDHm glioma models. **(D)** Quantification of [1-$^{13}$C]-glutamate production from [1-$^{13}$C]-$\alpha$-KG in VOR-resistant (MGG119, GSC580, NCH1681, NCH612) cells expressing a non-targeted sgRNA or PHGDH-targeted sgRNA and treated with vehicle or VOR for 72 h. **(E)** Quantification of steady-state D-2HG concentration in VOR-resistant (MGG119, GSC580, NCH1681, NCH612) cells expressing a non-targeted sgRNA or PHGDH-targeted sgRNA and treated with vehicle or VOR for 72 h. **(F)** Verification of PHGDH overexpression in VOR-sensitive (BT257, BT142, SF10417, TS603) IDHm glioma cells expressing an empty vector (EV) or PHGDH. **(G)** Quantification of [1-$^{13}$C]-glutamate production from [1-$^{13}$C]-$\alpha$-KG in VOR-sensitive (BT257, BT142, SF10417, TS603) IDHm glioma cells expressing an empty vector (EV) or PHGDH. Cells were treated with vehicle or VOR for 72 h. All studies were $n \geq 3$. Data is presented as mean ± standard deviation. * indicates statistical significance with $p<0.05$, ** indicates $p<0.01$, *** indicates $p<0.001$, **** indicates $p<0.0001$, ns is not significant.

Supplementary Figure 3

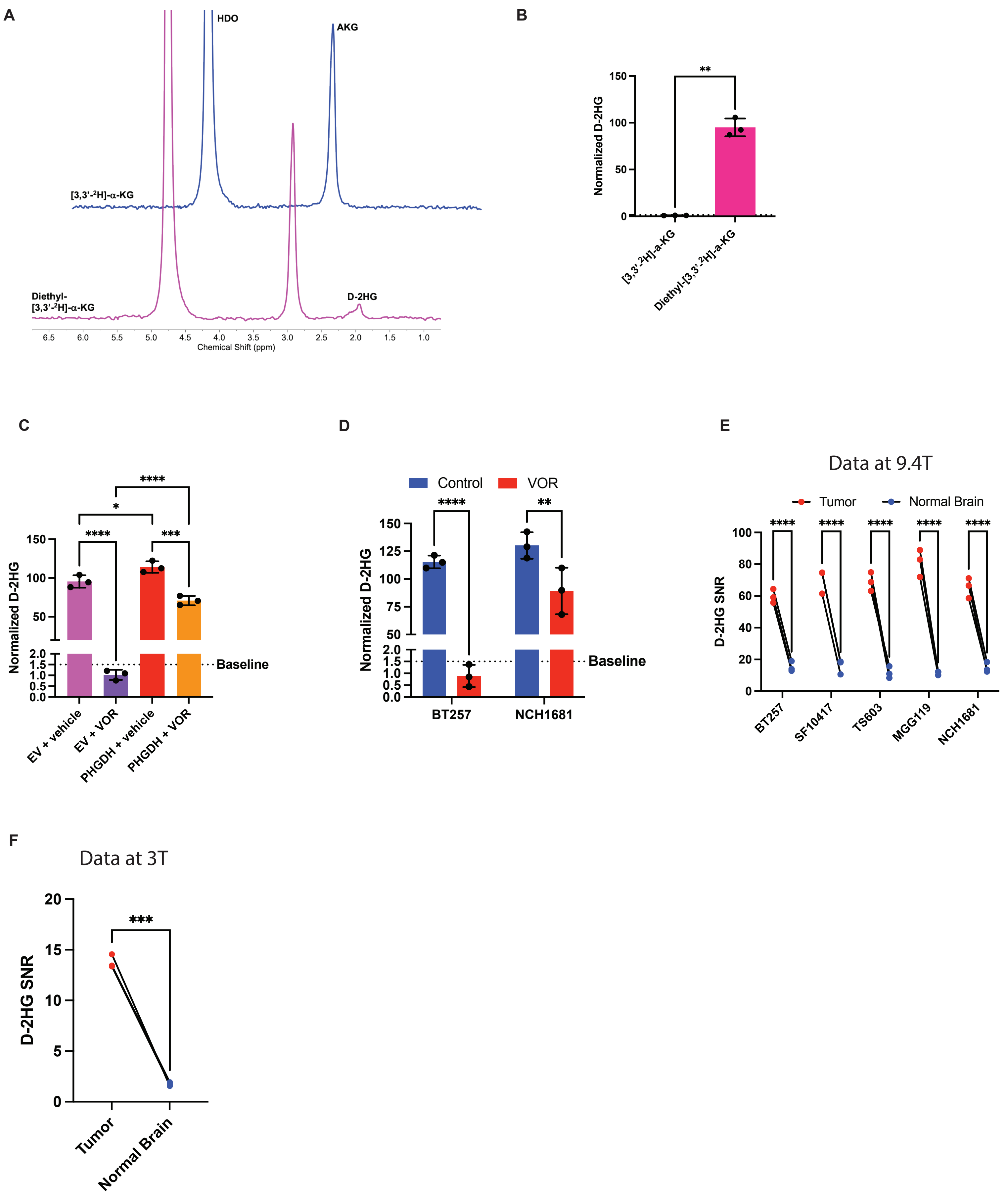

**Supplementary Figure 3. Diethyl-[3,3'-$^2$H]-$\alpha$-KG metabolism to D-2HG reports on tumor burden in IDHm gliomas. (A)** Representative $^2$H-MR spectra from NPAmut cells incubated with diethyl-[3,3'-$^2$H]-$\alpha$-KG (esterified) or [3,3'-$^2$H]-$\alpha$-KG (non-esterified) for 72 h. **(B)** Quantification of D-2HG production from diethyl-[3,3'-$^2$H]-$\alpha$-KG or [3,3'-$^2$H]-$\alpha$-KG in NPAmut cells. **(C)** Quantification of $^2$H-D-2HG produced from diethyl-[3,3'-$^2$H]-$\alpha$-KG in NPAmut cells overexpressing an EV ($NPAmut_{EV}$) or PHGDH ($NPAmut_{PHGDH}$). Cells were treated with vehicle or VOR for 72 h. **(D)** Quantification of $^2$H-D-2HG produced from diethyl-[3,3'-$^2$H]-$\alpha$-KG in patient-derived VOR-sensitive (BT257) or VOR-resistant (NCH1681) cells treated with vehicle or VOR for 72 h. **(E)** Quantification of the SNR of $^2$H-D-2HG produced from diethyl-[3,3'-$^2$H]-$\alpha$-KG in the tumor vs. the contralateral normal brain in mice bearing intracranial BT257, SF10417, TS603, MGG119, or NCH1681 tumors at 9.4T. **(F)** Quantification of the SNR of $^2$H-D-2HG produced from diethyl-[3,3'-$^2$H]-$\alpha$-KG in the tumor vs. the contralateral normal brain in mice bearing intracranial NPAmut tumors at 3T. All studies were $n \geq 3$. Data is presented as mean ± standard deviation. * indicates statistical significance with $p<0.05$, ** indicates $p<0.01$, *** indicates $p<0.001$, **** indicates $p<0.0001$, ns is not significant.

**Supplementary Figure 4**

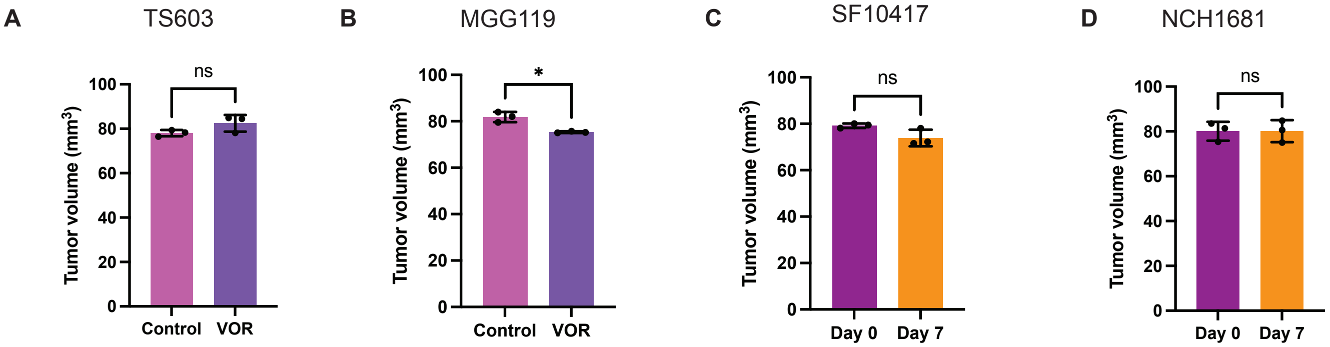

**Supplementary Figure 4. Diethyl-[3,3'-$^{2}$H]-$\alpha$-KG provides an early readout of treatment response that is independent of anatomical imaging.** Effect of treatment with VOR on tumor volume in mice bearing intracranial TS603 **(A)**, MGG119 **(B)**, SF10417 **(C)**, and NCH1681 **(D)** tumors. All studies were $n \geq 3$. Data is presented as mean ± standard deviation. * indicates statistical significance with $p<0.05$, ** indicates $p<0.01$, *** indicates $p<0.001$, **** indicates $p<0.0001$, ns is not significant.

**Table S1. Clinical information of IDHm inhibitor-treated patients included in this study.**

| Sample ID | IDH Mutation Status | Gender | Age | Diagnosis | Date of surgery | IDH Inhibitor treatment | IDHm inhibitor | Start date of IDHm inhibitor treatment | Progression | End date of IDHm inhibitor treatment | Date of tissue acquisition | Time to next intervention | Next intervention | Date of last followup | Vital Status |
|---|---|---|---|---|---|---|---|---|---|---|---|---|---|---|---|
| SF14058 | R132H | M | 44 | Anaplastic Astrocytoma, IDH-mutant | Jan-23 | Yes | IVO | Oct-21 | Yes | Jan-23 | Jan-25 | 15 months | Surgery at UCSF confirmed grade 3 | Jul-25 | Alive |
| SF12264 | R132H | M | 34 | Diffuse Astrocytoma, IDH-mutant | Apr-19 | Yes | VOR | Mar-19 | Yes | Feb-24 | Apr-25 | 59 months | Surgery at UCSF confirmed grade 3 | May-25 | Alive |
| SF12239 | R132H | F | 35 | Anaplastic Oligodendroglioma, IDH-mutant, 1p19q code | Mar-19 | Yes | IVO | Aug-18 | Yes | Mar-19 | Mar-19 | 7 months | Surgery at UCSF confirmed grade 3 | Sep-20 | Dead |
| SF12098 | R132H | M | 38 | Oligodendroglioma, IDH-mutant, 1p19q codeleted | Dec-18 | Yes | IVO | Nov-18 | Yes | Apr-22 | Dec-18 | 40 months | Surgery at MSKCC confirmed recurrent grade 2 | Jul-25 | Alive |
| SF12819 | R132H | F | 21 | Anaplastic Astrocytoma, IDH-mutant | Jun-20 | Yes | VOR | Jun-17 | Yes | May-18 | Jun-20 | 11 months | Proton radiation | Mar-21 | Dead |
| SF13345 | R132H | M | 32 | Oligodendroglioma, IDH-mutant, 1p19q codeleted | Jul-21 | Yes | IVO | Dec-20 | Yes | Oct-24 | Jul-21 | 46 months | Surgery at UCSF confirmed recurrent grade 2 | Jul-25 | Alive |

**Table S2. Clinical information of untreated external control patients included in this study.**

| Sample ID | IDH Mutation Status | Gender | Age | Diagnosis | Date of surgery | IDH Inhibitor treatment |
|---|---|---|---|---|---|---|
| SF11698 | R132H | M | 47 | Anaplastic Astrocytoma, IDH-mutant | Mar-18 | No |
| SF11317 | R132H | F | 30 | Diffuse Astrocytoma, IDH-mutant | Jun-17 | No |
| SF13091 | R132H | M | 49 | Anaplastic Oligodendroglioma, IDH-mutant, 1p19q code | Jan-21 | No |
| SF14089 | R132H | M | 47 | Oligodendroglioma, IDH-mutant, 1p19q codeleted | Feb-23 | No |
| SF12853 | R132H | M | 39 | GBM, IDH-mutant | Aug-20 | No |
| SF13081 | R132H | M | 36 | Diffuse Astrocytoma, IDH-mutant | Jan-21 | No |
| SF13062 | R132H | M | 59 | Oligodendroglioma, IDH-mutant, 1p19q codeleted | Jan-21 | No |
